# Molecular biogeography and host relations of a parasitoid fly

**DOI:** 10.1101/576892

**Authors:** David A. Gray, Henry D. Kunerth, Marlene Zuk, William H. Cade, Susan L. Balenger

## Abstract

Successful geographic range expansion by parasites and parasitoids may also require host range expansion. Thus the evolutionary advantages of host specialization may trade off against the ability to exploit new host species encountered in new geographic regions. Here we use molecular techniques and confirmed host records to examine biogeography, population divergence, and host flexibility of the parasitoid fly, *Ormia ochracea* (Bigot). Gravid females of this fly find their cricket hosts acoustically by eavesdropping on male cricket calling songs; these songs vary greatly among the known host species of crickets. Using both nuclear and mitochondrial genetic markers, we (1) describe the geographical distribution and sub-division of genetic variation in *O. ochracea* from across the continental United States, the Mexican states of Sonora and Oaxaca, and populations introduced to Hawaii; (2) demonstrate that the distribution of genetic variation among fly populations is consistent with a single widespread species with regional host specialization, rather than locally differentiated cryptic species, (3) identify the more-probable source populations for the flies introduced to the Hawaiian islands; (4) examine genetic variation and sub-structure within Hawaii; and (5) discuss specialization and lability in host-finding behavior in light of the diversity of cricket songs serving as host cues in different geographically separate populations.

## Introduction

Evolutionary specialization is often viewed as a double-edged sword: specialization may facilitate efficient exploitation of favored resources, but may also inhibit exploitation of novel resources. Specialization has often been viewed as an evolutionary ‘dead-end’ (Raia and Fortelius, 2013, Jaenike, 1990, Kelley and Farrell, 1998), although recent research has revealed considerable flexibility among specialist lineages and occasional ‘reversals’ from specialized to more generalized niches (Vamosi et al., 2014, Gompert et al., 2015). The retention of evolutionary lability may be especially relevant for geographic range expansion; indeed ‘generalist’ species are often among the most invasive (Romanuk et al., 2009) – a pattern found among plants, arthropods, mammals and birds (Higgins and Richardson, 2014, González-Suárez et al., 2015, Blackburn and Duncan, 2001, Snyder and Evans, 2006). For specialist species to expand their geographic range, they must readily encounter suitable resources, exhibit phenotypic plasticity enabling adoption of novel resources, and/or show rapid evolutionary adaptation.

Parasitoid insects, especially Ichneumonid and Braconid wasps (Hymenoptera) and Tachinid flies (Diptera), are especially illuminating for studies of host specialization, ranging from extreme generalists to extreme specialists (Quicke, 2014, Stireman et al., 2006). Some species are sufficiently host specific to be used for classical biological control of pests (Parkman et al., 1996, Vargas et al., 2007), others routinely utilize a broad range of hosts (Stireman, 2005, Tschorsnig, 2017, Arnaud, 1978), and in other cases, presumed generalists are later revealed to be complexes of cryptic specialists (Smith et al., 2008).

Within the ca. 9000 species of Tachinids, the Ormiini tribe represents a small group (ca. 68 described species) of highly specialized flies (Sabrosky, 1953a, Sabrosky, 1953b, Lehmann, 2003). Several specializations are noteworthy for the entire group (so far as is known): all are parasitoids of crickets or katydids (Ensifera, Orthoptera); all locate their (principally male) hosts using a specialized ear (Edgecomb et al., 1995, Hedwig and Robert, 2014) to eavesdrop on their male host’s mating song (Lehmann, 2003, Cade, 1975, Allen, 1995); all have sclerotized planidiform larvae which are somewhat mobile and actively burrow into the host (Cantrell, 1988, Adamo et al., 1995b). Within this group, all genera with known hosts parasitize katydids (Tettigoniidae); in the genus *Ormia* most species parasitize katydids but three species attack crickets and mole crickets (Gryllidae and Gryllotalpidae) (Lehmann, 2003). The shift from katydids to crickets and mole crickets represents a significant shift in female fly hearing towards lower frequency sounds (ca. 4-5 kHz in crickets and ca. 2-3 kHz in mole crickets) than are typical of most katydids (often >>10kHz). Utilization of katydids with relatively low frequency calls may have facilitated the evolutionary transition to crickets and mole crickets. For example, certain katydid hosts of Ormiines have relatively low frequency calls, e.g. ca. 5-6 kHz in *Sciarasaga quadrata* (host of *Homotrixa alleni*) (Allen et al., 1999); ca. 7 kHz in *Neoconocephalus robustus* (host of *O. brevicornis*) (Nutting, 1953); ca. 8 kHz in *Orchelimum pulchellum* (one of several hosts of *O. lineifrons*) (Shapiro, 1995).

Within *Ormia, O. ochracea* has been most extensively studied. Peak sensitivity of female hearing closely matches or is at slightly higher frequencies than typical male calling song (Robert et al., 1992). The current geographic range attributed to this species extends from Florida (Walker and Wineriter, 1991), across the southern Gulf States (Henne and Johnson, 2001), into Texas (Cade, 1975), Arizona (Sakaguchi and Gray, 2011), California (Wagner, 1996), and Mexico (Sabrosky, 1953b); throughout this range it parasitizes various species of *Gryllus* field crickets (see below). In addition, *O. ochracea* was introduced to Hawaii by at least 1989 (Evenhuis, 2003), where it parasitizes *Teleogryllus oceanicus,* itself introduced to Hawaii by at least 1877 (Kevan, 1990) and possibly earlier, perhaps facilitated by Polynesian settlement (Tinghitella et al., 2011). Localized populations of *O. ochracea* show varying degrees of host specialization: flies in Florida almost exclusively parasitize *Gryllus rubens* (Walker, 1993, Walker and Wineriter, 1991); flies in Texas primarily parasitize *G. texensis* (Cade, 1975); flies in Arizona regularly parasitize multiple *Gryllus* species (Sakaguchi and Gray, 2011); flies in southern California primarily parasitize *G. lineaticeps* (Wagner, 1996, Wagner and Basolo, 2007); as noted above, Hawaiian flies parasitize *T. oceanicus*. Remarkably, playback experiments in Florida, Texas, California, and Hawaii, which simultaneously presented the songs of *G. rubens, G. texensis, G. lineaticeps*, and *T. oceanicus*, revealed that each fly population showed a significant (but not exclusive) preference for the song of its primary local host species of cricket (Gray et al., 2007). This suggests an even further degree of host specialization in these flies – possibly indicative of cryptic host races or species as has been found in other Tachinids (Smith et al., 2008, Smith et al., 2006). Determining the extent to which geographic and host range subdivision is coupled with genetic subdivision is thus one of the goals of this study.

Successful establishment of *O. ochracea* in Hawaii represents a significant expansion of both the geographic and host range of the fly. How can such a specialist invade, switch to a novel host with a strongly divergent song structure, and in the course of a few decades come to prefer that novel host’s song to the songs of ancestral hosts? Two of our aims in this paper are to use mitochondrial and nuclear markers both to examine genetic variation within Hawaii and to identify the more-likely continental source population(s) of those Hawaiian flies, and thereby the most likely types of recent ancestral host songs. This necessitates broad sampling of continental populations, and we therefore expand upon the previous work in the USA and include flies from populations in both northern and southern Mexico, as well as catalog the confirmed host species and their songs in each of these areas. We apply standard phylogeographic analyses to mitochondrial DNA sequence data, including outgroup species of *Ormia*, and we adopt a population genetic approach to analysis of microsatellite nuclear markers.

## Methods

### Fly collection

We collected flies at mesh screen and/or bottle traps using playbacks of cricket songs (Walker, 1989); we also collected a small number of flies at lights or as they emerged from field-collected crickets. Table 1 provides details of locations and dates of sampling. Collected flies were preserved in ethanol until DNA extraction and further analysis. We extracted DNA using a Qiagen DNeasy tissue kit according to the manufacturer’s instructions. We used entire flies as source tissue for all of the mainland and 13 of the Hawaiian flies, and head and thorax tissue for the remainder of the Hawaiian flies. In theory, the whole tissue extractions could include DNA from larvae, although the amounts of such DNA would be trivial compared to maternal DNA. We quantified DNA using a Nanodrop system and adjusted concentrations to between 20 and 75 ng/ul.

**Table 1.** Sample collection data; not all specimens were used in all analyses.

| **Region** | **Locality** | **Dates** | **N** | **Collector(s)** |
| --- | --- | --- | --- | --- |
| **Florida** | Gainesville, FL | Aug. 2002 | 41 | DAG |
| **Texas** | San Antonio, TX | Sept. 2002 | 5 | WHC |
|  | Austin, TX | Sept. 2002, 2004 | 29 | WHC & S. Walker 2002; DAG 2004 |
|  | Huntsville, TX | Sept. 2002 | 1 | S. Walker |
| **Arizona** | Sedona, AZ | Aug. 2004 | 12 | DAG |
|  | Oak Creek, AZ | Aug. 2004 | 6 | DAG |
|  | Holbrook, AZ | Aug. 2002 | 1 | DAG |
|  | Verde River, AZ | Aug. 2004 | 3 | DAG |
|  | Madera Canyon, AZ | Aug. 2004 | 10 | DAG |
|  | KOFA, AZ | Sept. 2005 | 2 | DAG |
|  | Yuma, AZ | Nov. 2003 | 2 | A. Izzo |
|  | Parker Canyon, AZ | Aug. 2004 | 2 | DAG |
|  | Petroglyph, AZ | Sept. 2006 | 16 | DAG |
|  | Pinery Canyon, AZ | Sept. 2004 | 5 | DAG |
|  | Portal, AZ | Aug. 2003 | 1 | DAG |
| **Sonora** | Alamos, Sonora, MX | July 2006 | 17 | DAG |
| **Oaxaca** | San Pablo Etla, Oaxaca, MX | Nov. 2014 | 13 | DAG |
| **California** | Malibu Creek, CA | Sept. & Oct. 2003, 2004 | 22 | DAG |
|  | Stunt Ranch, CA | Sept. 2002 | 10 | DAG |
|  | Santa Margarita Reserve, CA | Sept. 2003 | 5 | DAG |
| **Hawaii** | Kauai, HI | Feb. & Aug. 2014 | 24 | MZ & SLB |
|  | Hilo, HI | Mar. 2003; Feb. & Aug. 2014 | 33 | WHC 2003; MZ & SLB 2014 |
|  | Oahu, HI | Feb. 2014 | 4 | MZ & SLB |
| **Outgroups** |  |  |  |  |
| *Ormia depleta* | Gainesville, FL | Dec. 2003 | 2 | H. Frank, via T. J. Walker |
| *Ormia lineifrons* | Gainesville, FL | Dec. 2003 | 2 | H. Frank, via T. J. Walker |

### Genetic Markers & Analysis

We analyzed population structure using both mitochondrial and nuclear markers. For mtDNA, we analyzed a section of *Cytochrome C Oxidase subunit I* (hereafter COI) PCR amplified in two overlapping fragments with ‘universal’ primer pairs Jerry-Pat and Ron-Nancy (Simon et al., 1994), resulting in 1111 bp after alignment. In addition, we developed nuclear microsatellite markers *de novo* for this project. Marker discovery was performed by 454 sequencing at the Cornell University Life Sciences Core Laboratories Center with further validation done by SLB and HDK. We identified and tested 17 msat markers from this dataset consisting of 3, 4, and 6 bp repeats. PCR conditions followed a ‘touchdown’ protocol of 95° for 40 seconds, 66° for 45 seconds, and 72° for 45 seconds. The annealing step was reduced by one degree every cycle for the first seven cycles. Cycles 8-35 followed a pattern of 95° for 40s, 58° for 45s, and 72° for 45s. PCR products were stored at −20°C until genotyped. Individuals were genotyped at microsatellite loci by the University of Minnesota Genomics Center on an Applied Biosystems 3730xl DNA Analyzer. We scored alleles for fragment size manually using Peak Scanner 2.0 software. Multiple independent analysts scored the same products to assure veracity of the calls. If no clear designation could be made or alleles did not amplify, we scored the data as missing.

### Bioinformatic Analyses

Prior to analysis of microsatellite fragments, we filtered individuals and loci for missing data. A strict cutoff of >25% missing data led to the exclusion of 6 loci. Following this filter, we excluded any individuals with missing data at 3 or more loci. The final dataset included 274 individuals genotyped at 11 loci with between 6 and 17 alleles per locus (Table 2). To estimate the number of alleles and private alleles accurately given unequal sample sizes per population, we performed a rarefaction analysis using HP-Rare (Kalinowski, 2005). We visualized population genetic variation using a discriminant function analysis of principal components (DAPC) with 80 principal components and 4 discriminant functions using the adegenet (Jombart, 2008, Jombart and Ahmed, 2011) and pegas (Paradis, 2010) packages in R.

**Table 2.**
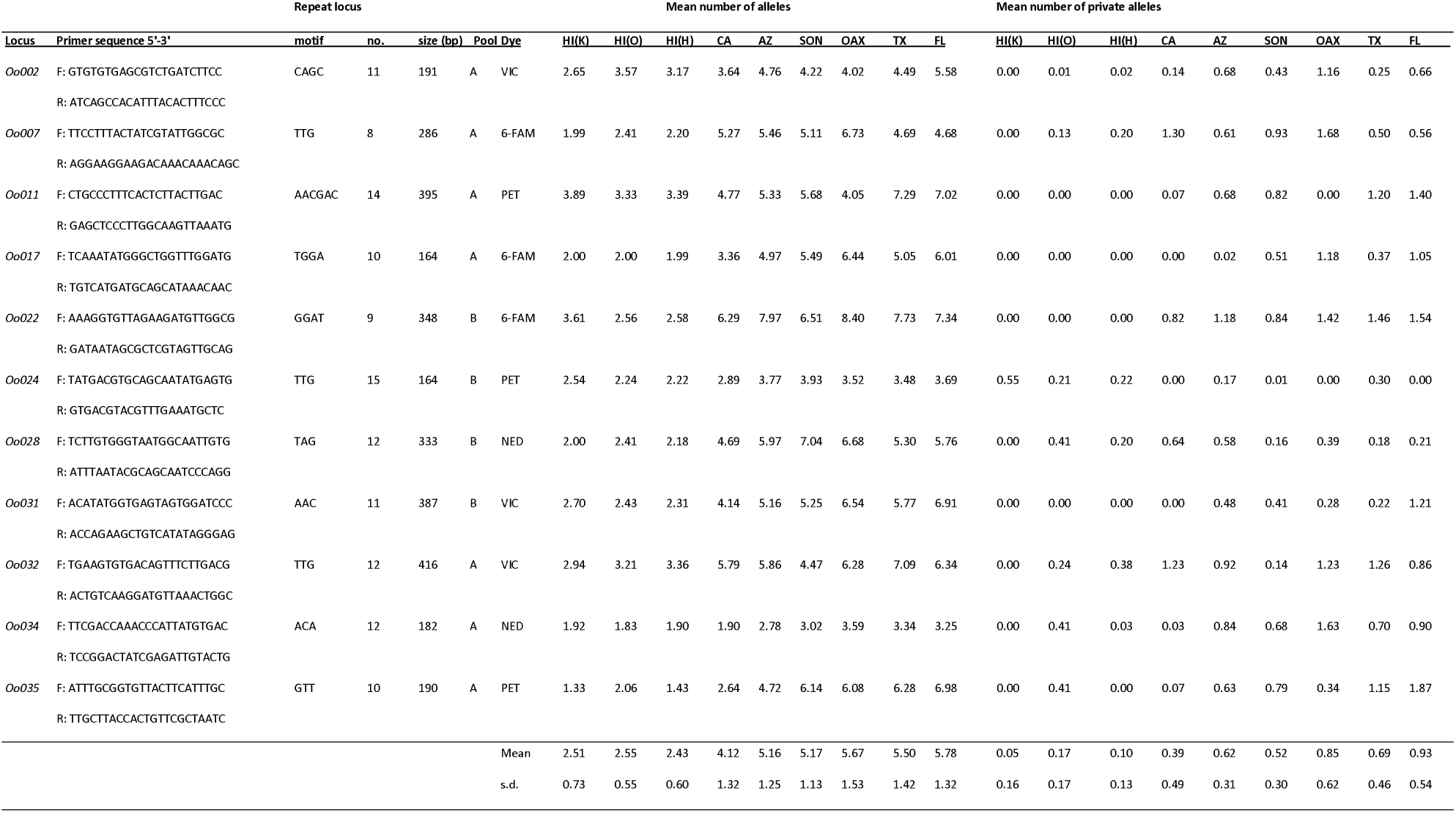
Locus primer and allelic richness statistics

To visualize genetic structure, we implemented the Bayesian analysis program STRUCTURE v2.3.4 using an admixture model with correlated allele frequencies. We used a burn-in of 50,000 steps and 100,000 MCMC iterations. We conducted separate runs for the full dataset, a dataset with the Hawaiian samples excluded, and a dataset of only Hawaiian samples. For the full dataset, we performed 5 runs each for k = 2-9. To infer the likely number of genetic clusters, we used both the Ln estimated probability of the data from STRUCTURE and the Evanno method utilizing Δk (Evanno et al., 2005).

We calculated pairwise estimates of Fst (Weir and Cockerham, 1984) and Nei’s genetic distance between populations using the R packages adegenet and ade4 (Chessel et al., 2004), and we calculated expected and observed heterozygosity using adegenet.

We built a mitochondrial haplotype network using 55 haplotypes from 1111 bp of COI sequences from 275 individuals using the R package pegas (Paradis, 2010) with default parameters.

### Host Ranges & Songs

To provide context for understanding the degree of host specialization, we present in this paper the songs of confirmed hosts in each of the geographic regions studied. We present only hosts confirmed to be naturally parasitized by development of *O. ochracea* from field-collected crickets. We suspect that a few additional host species will be confirmed in the USA, especially if the species is only occasionally parasitized, and we expect that many more species are parasitized in southern and central Mexico; this reflects the status of current knowledge of *Gryllus* systematics and the extent of field sampling. Many of the confirmed host species are not yet officially described (DB Weissman and DA Gray, in prep.); to provide continuity within the literature we use provisional manuscript names here and note that the names are disclaimed as unavailable per Article 8.3 of the ICZN.

In an attempt to quantify relative song differences, we created a Euclidean song distance matrix using *matrix <-dist(songdata)* function in R. Song variables were: dominant frequency (kHz), pulse rate, pulses per chirp or trill (ln transformed), pulse duty cycle, song type (chirp, trill, stutter-trill, complex stutter-trill), chirps per trill (for stutter-trillers), as well as introductory pulses per trill and introductory pulse rate (for complex stutter-trillers). Prior to matrix calculation, the raw song data were normalized as z-scores. The resulting song distance matrix has the advantage of objectively showing unit-less quantitative differences among species, but has the disadvantage that the different song features are not weighted by their perceptual importance to *O. ochracea*, which would be preferable but is not currently possible.

## Results

### Nuclear and mitochondrial genetics

Following filtration at missing data cutoffs, 274 individuals and 11 loci were included in the final msat dataset, with 1.86% data missing. Heterozygosity across all individuals was 50.9%. The Hawaiian populations showed a drastic decrease in heterozygosity (Table 3). The rarefaction analysis also suggested a substantial decrease in both total and private allelic diversity within the Hawaiian populations (Table 2).

**Table 3.** Population sample sizes and heterozygosity for nuclear msat loci.

| Population | Sample size | N. alleles | Heterozygosity<br>(expected) | Heterozygosity<br>(observed) |
| --- | --- | --- | --- | --- |
| Kauai | 20 | 29 | 0.437 | 0.367 |
| Oahu | 28 | 31 | 0.438 | 0.367 |
| Hilo | 32 | 34 | 0.401 | 0.321 |
| California | 32 | 62 | 0.588 | 0.478 |
| Arizona | 57 | 95 | 0.667 | 0.612 |
| Sonora | 17 | 70 | 0.677 | 0.588 |
| Oaxaca | 13 | 70 | 0.724 | 0.607 |
| Texas | 35 | 91 | 0.714 | 0.604 |
| Florida | 40 | 95 | 0.741 | 0.638 |

Analysis of Nei’s genetic distances documented a clear split between Hawaiian and mainland populations (Table 4), with Hawaiian populations more similar to western mainland populations. Longitude explained the primary axis of variation among the mainland populations, with a clear east-west gradient evident in both the DAPC and mtDNA haplotype network (Fig. 1), as well as in the pairwise Fst and Nei’s distances (Table 4).

**Table 4.** Pairwise F_ST_ (above diagonal) and Nei’s genetic distance (below diagonal) by population.

|  | **Kauai** | **Oahu** | **Hilo** | **California** | **Arizona** | **Sonora** | **Oaxaca** | **Texas** | **Florida** |
| --- | --- | --- | --- | --- | --- | --- | --- | --- | --- |
| **Kauai** | - | 0.027 | 0.057 | 0.092 | 0.071 | 0.105 | 0.109 | 0.091 | 0.087 |
| **Oahu** | 0.044 | - | 0.047 | 0.088 | 0.079 | 0.099 | 0.095 | 0.100 | 0.098 |
| **Hilo** | 0.096 | 0.073 | - | 0.114 | 0.097 | 0.124 | 0.118 | 0.127 | 0.122 |
| **California** | 0.263 | 0.229 | 0.279 | - | 0.024 | 0.034 | 0.049 | 0.055 | 0.060 |
| **Arizona** | 0.282 | 0.267 | 0.291 | 0.088 | - | 0.011 | 0.019 | 0.031 | 0.035 |
| **Sonora** | 0.290 | 0.286 | 0.344 | 0.127 | 0.067 | - | 0.032 | 0.026 | 0.026 |
| **Oaxaca** | 0.327 | 0.305 | 0.365 | 0.235 | 0.151 | 0.169 | - | 0.022 | 0.021 |
| **Texas** | 0.331 | 0.332 | 0.394 | 0.231 | 0.165 | 0.149 | 0.158 | - | 0.008 |
| **Florida** | 0.337 | 0.336 | 0.388 | 0.273 | 0.187 | 0.167 | 0.171 | 0.045 | - |

**Figure 1.**
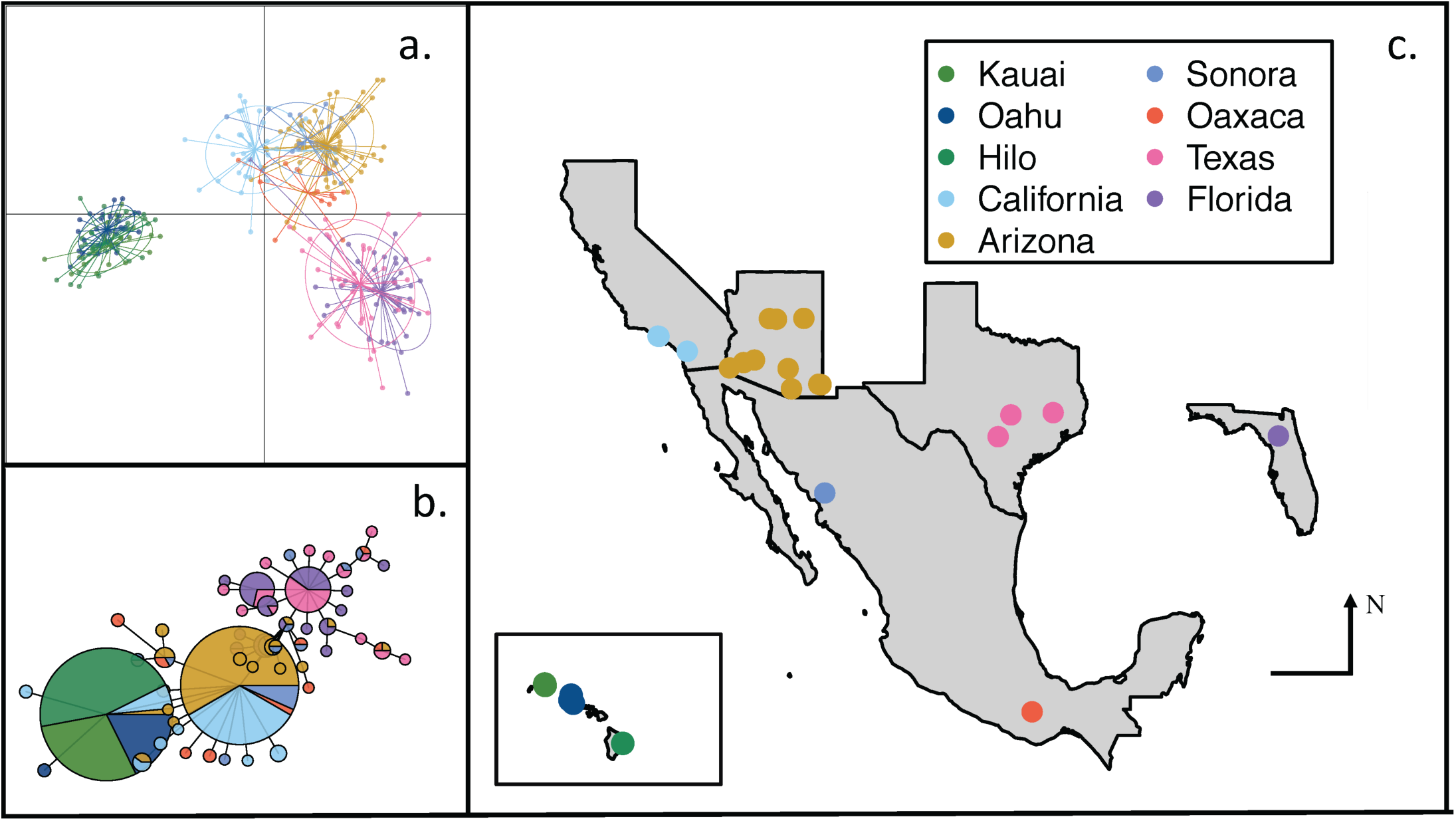
a) DAPC clustering analysis. Individuals are marked as points with ellipses representing 75% of ed data. b) Haplotype network of 55 haplotypes of 1111bp of mitochondrial COI gene c) Map of collection sites.

For the full dataset, STRUCTURE analyses indicated the strongest support for k=2 genetic clusters (Fig. 2) separating Hawaiian from mainland populations, however support for k=3 clusters was also high, which further divided the mainland populations into eastern and western subsets (Fig. 2). STRUCTURE plots for within Hawaii (k=2 and k=3) and mainland (k=2, k=3, and k=6) are in Supplemental Materials Figs. S1 and S2.

**Figure 2.**
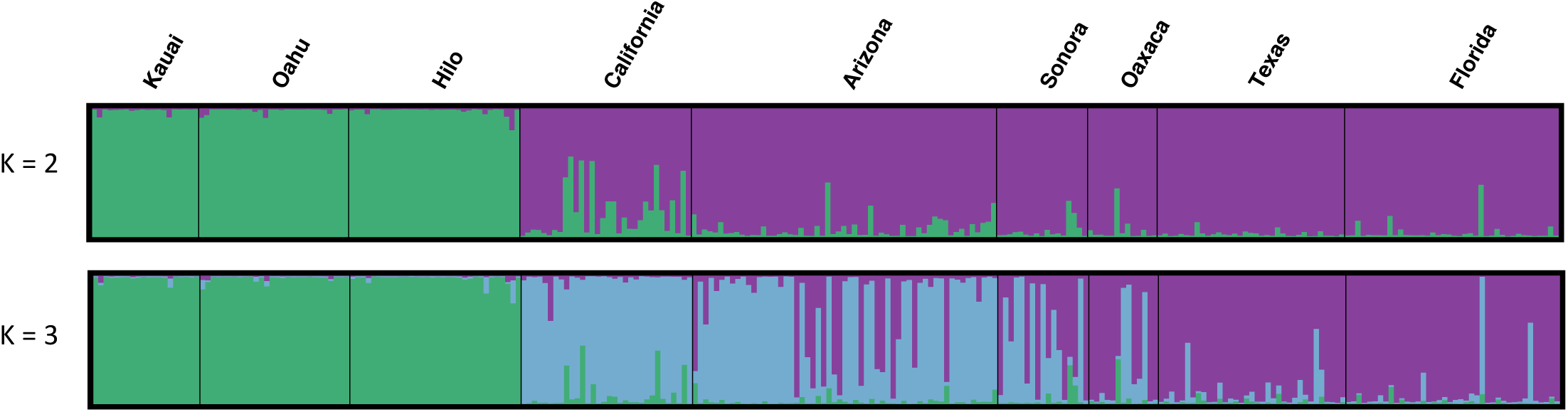
Bayesian clustering analysis implemented by STRUCTURE software (Pritchard et al. 2000). shows clustering into two genetic groups (K = 2) and the bottom panel shows clustering genetic groups (K = 3).

The mtDNA haplotype network (Fig. 1b) also showed (1) low genetic variation within Hawaii, (2) affinity of the Hawaiian sequences for the western mainland (i.e. California) sequences, and (3) a longitudinal geographic structure within the mainland populations. Oaxaca had a high diversity of haplotypes shared with all other mainland populations.

Given the apparent distinctness of the Hawaiian populations, it is important to emphasize that these patterns reflect founder effects, and concomitant change in allele frequency in Hawaii, not the development of novel genetic variation in Hawaii. This is most easily seen in allele frequency histograms which show that the Hawaiian genetic variation is effectively a simple subset of the genetic variation found in western mainland populations, themselves a simple subset of the genetic variation found in Florida, Texas, and Mexico populations (see Fig. 3 for a representative locus; figures for all other loci show similar patterns and are presented as Supplemental Materials Figures S3-S12).

**Figure 3.**
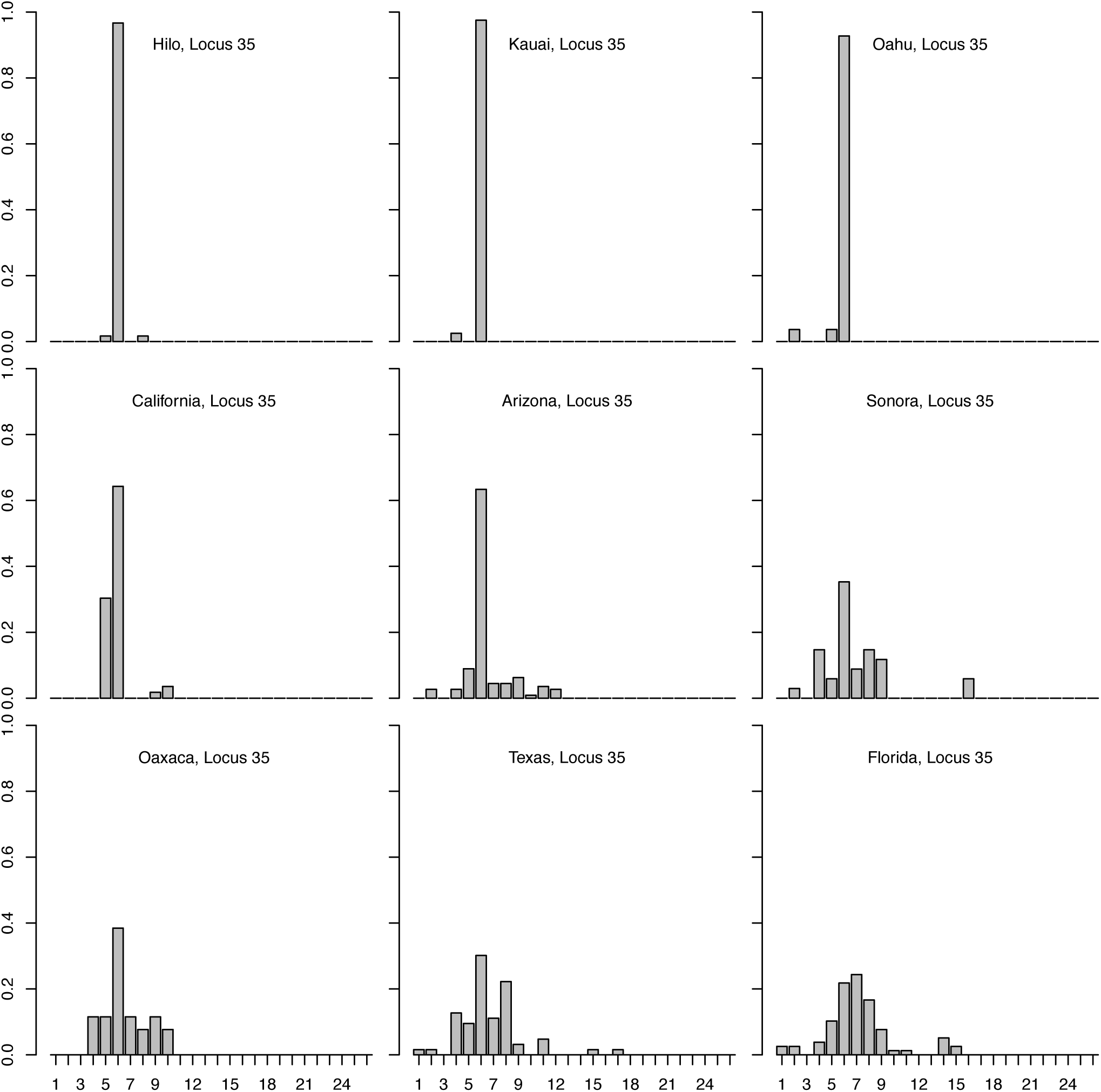
Allele frequency histograms for msat locus 35 for each population.

### Host range and song structures

Confirmed host species, geographic range information, as well as host calling song type, frequency, pulse rate, and pulses/chirp are presented in Table 5. Songs of confirmed host species vary dramatically, from simple chirps to complex trills; see waveform oscillograms and frequency spectrograms in Figures 4 and 5, respectively.

**Table 5.**
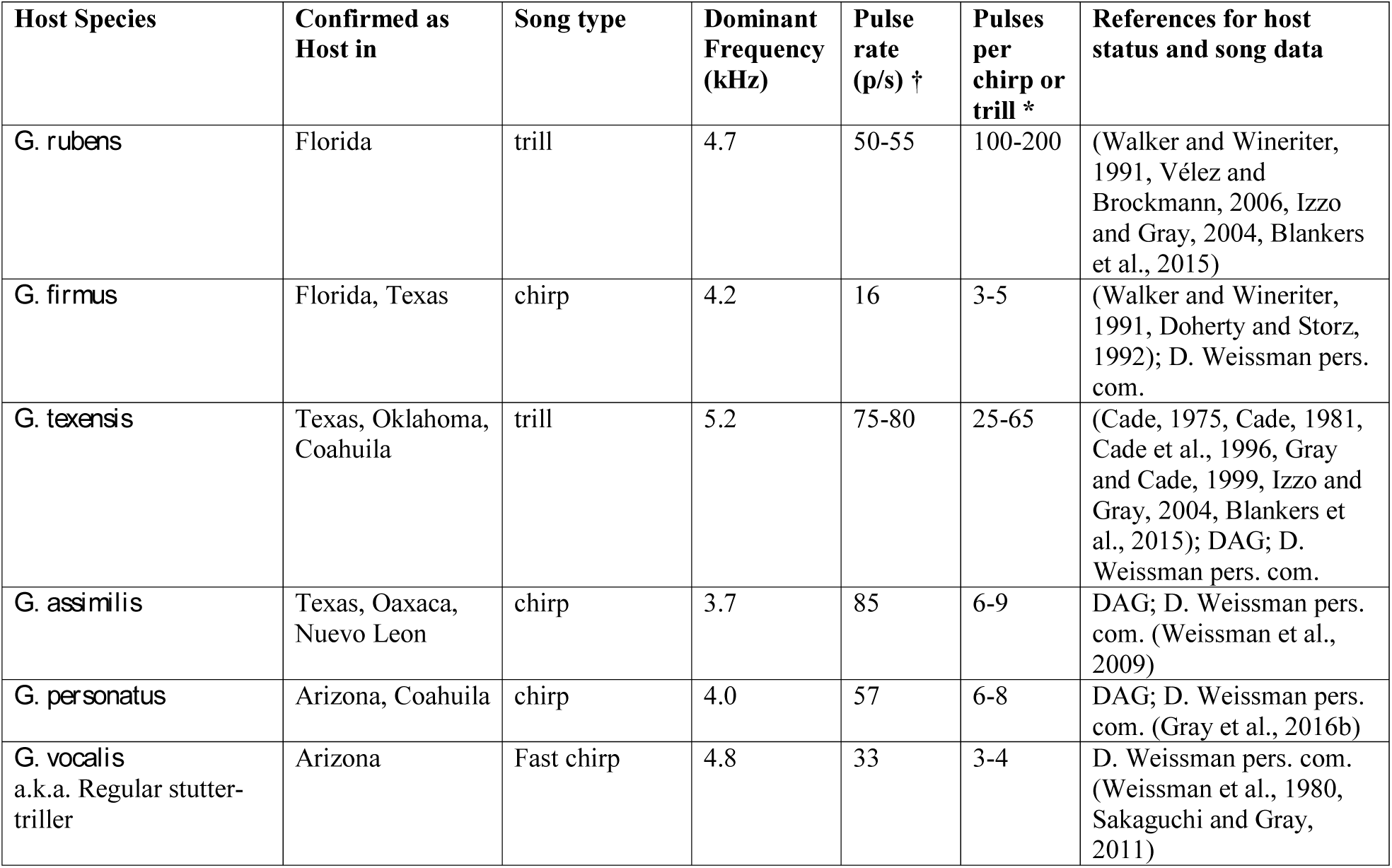
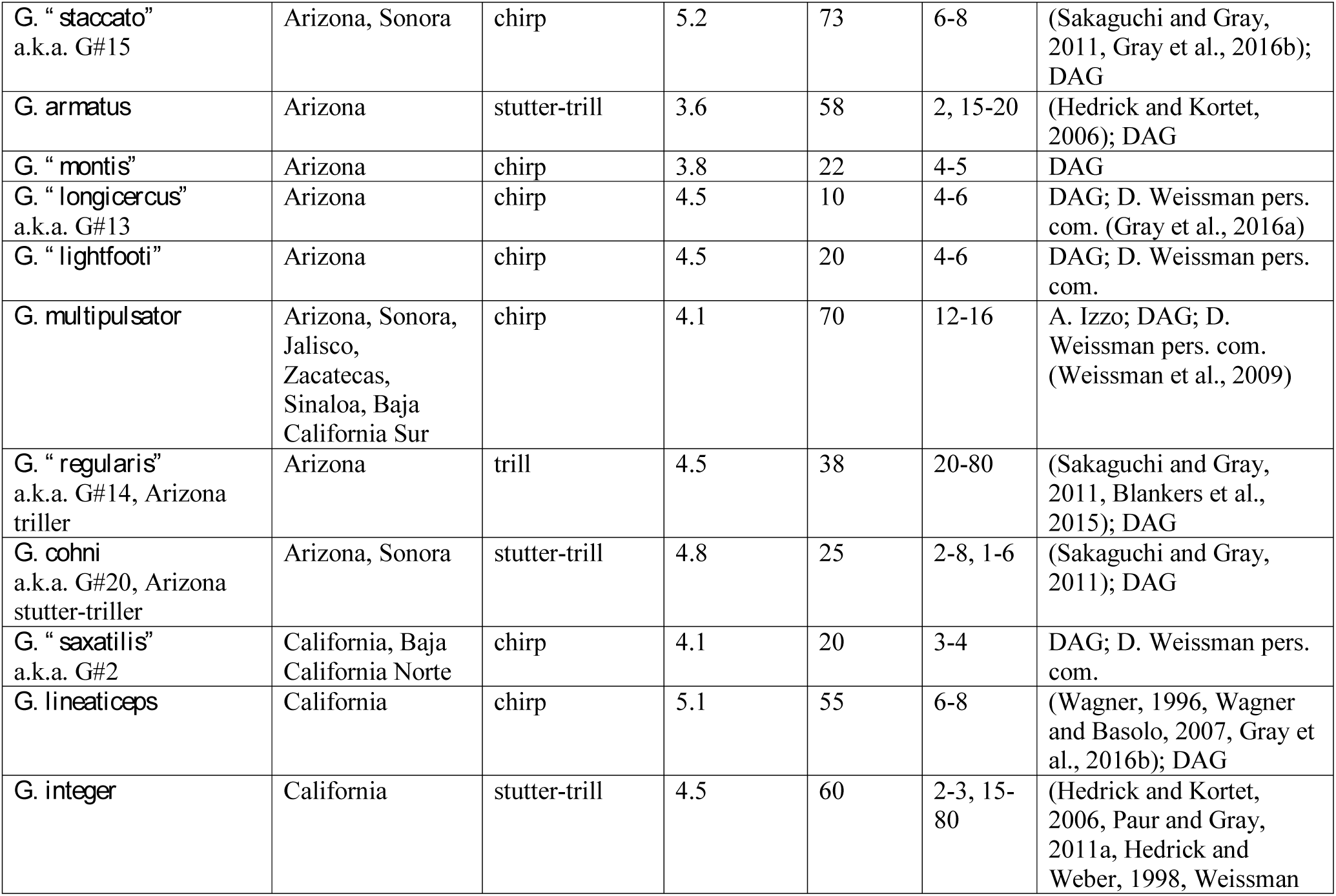
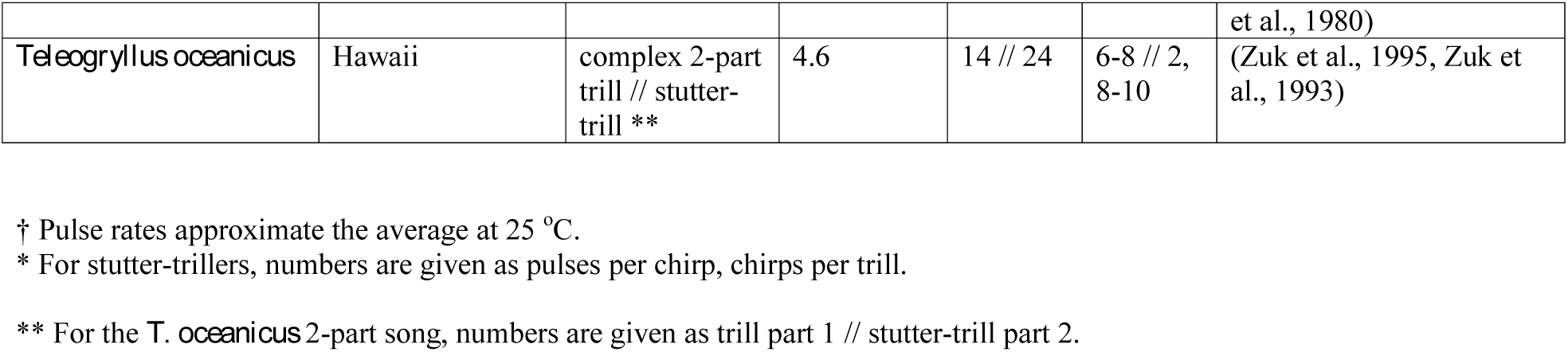
Confirmed hosts of *Ormia ochracea.*

| Host Species | Confirmed as Host in | Song type | Dominant Frequency (kHz) | Pulse rate (p/s) † | Pulses per chirp or trill * | References for host status and song data |
| --- | --- | --- | --- | --- | --- | --- |
| *G. rubens* | Florida | trill | 4.7 | 50-55 | 100-200 | (Walker and Wineriter, 1991, Vélez and Brockmann, 2006, Izzo and Gray, 2004, Blankers et al., 2015) |
| *G. firmus* | Florida, Texas | chirp | 4.2 | 16 | 3-5 | (Walker and Wineriter, 1991, Doherty and Storz, 1992); D. Weissman pers. com. |
| *G. texensis* | Texas, Oklahoma, Coahuila | trill | 5.2 | 75-80 | 25-65 | (Cade, 1975, Cade, 1981, Cade et al., 1996, Gray and Cade, 1999, Izzo and Gray, 2004, Blankers et al., 2015); DAG; D. Weissman pers. com. |
| *G. assimilis* | Texas, Oaxaca, Nuevo Leon | chirp | 3.7 | 85 | 6-9 | DAG; D. Weissman pers. com. (Weissman et al., 2009) |
| *G. personatus* | Arizona, Coahuila | chirp | 4.0 | 57 | 6-8 | DAG; D. Weissman pers. com. (Gray et al., 2016b) |
| *G. vocalis*<br>a.k.a. Regular stutter-triller | Arizona | Fast chirp | 4.8 | 33 | 3-4 | D. Weissman pers. com. (Weissman et al., 1980, Sakaguchi and Gray, 2011) |
| *G. "staccato"*<br>a.k.a. G#15 | Arizona, Sonora | chirp | 5.2 | 73 | 6-8 | (Sakaguchi and Gray, 2011, Gray et al., 2016b); DAG |
| *G. armatus* | Arizona | stutter-trill | 3.6 | 58 | 2, 15-20 | (Hedrick and Kortet, 2006); DAG |
| *G. "montis"* | Arizona | chirp | 3.8 | 22 | 4-5 | DAG |
| *G. "longicercus"*<br>a.k.a. G#13 | Arizona | chirp | 4.5 | 10 | 4-6 | DAG; D. Weissman pers. com. (Gray et al., 2016a) |
| *G. "lightfooti"* | Arizona | chirp | 4.5 | 20 | 4-6 | DAG; D. Weissman pers. com. |
| *G. multipulsator* | Arizona, Sonora, Jalisco, Zacatecas, Sinaloa, Baja California Sur | chirp | 4.1 | 70 | 12-16 | A. Izzo; DAG; D. Weissman pers. com. (Weissman et al., 2009) |
| *G. "regularis"*<br>a.k.a. G#14, Arizona triller | Arizona | trill | 4.5 | 38 | 20-80 | (Sakaguchi and Gray, 2011, Blankers et al., 2015); DAG |
| *G. cohni*<br>a.k.a. G#20, Arizona stutter-triller | Arizona, Sonora | stutter-trill | 4.8 | 25 | 2-8, 1-6 | (Sakaguchi and Gray, 2011); DAG |
| *G. "saxatilis"*<br>a.k.a. G#2 | California, Baja California Norte | chirp | 4.1 | 20 | 3-4 | DAG; D. Weissman pers. com. |
| *G. lineaticeps* | California | chirp | 5.1 | 55 | 6-8 | (Wagner, 1996, Wagner and Basolo, 2007, Gray et al., 2016b); DAG |
| *G. integer* | California | stutter-trill | 4.5 | 60 | 2-3, 15-80 | (Hedrick and Kortet, 2006, Paur and Gray, 2011a, Hedrick and Weber, 1998, Weissman et al., 1980) |
| *Teleogryllus oceanicus* | Hawaii | complex 2-part<br>trill // stutter-trill ** | 4.6 | 14 // 24 | 6-8 // 2,<br>8-10 | (Zuk et al., 1995, Zuk et<br>al., 1993) |

† Pulse rates approximate the average at 25 °C.
\* For stutter-trillers, numbers are given as pulses per chirp, chirps per trill.
\*\* For the *T. oceanicus* 2-part song, numbers are given as trill part 1 // stutter-trill part 2.

**Figure 4.**
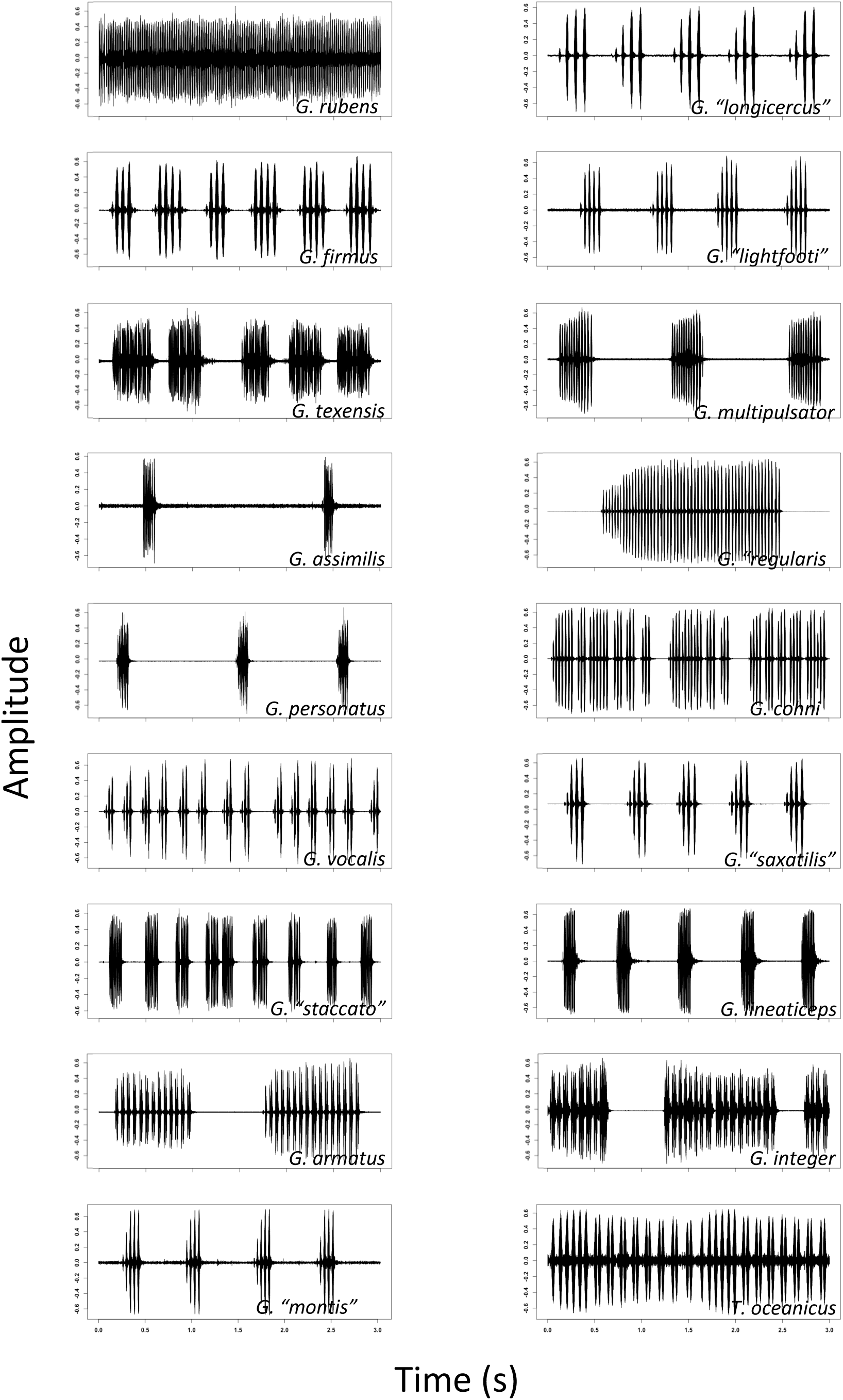
Waveform oscillograms of 3 seconds of song from confirmed host species showing overall song structure(chirps/trills).

**Figure 5.**
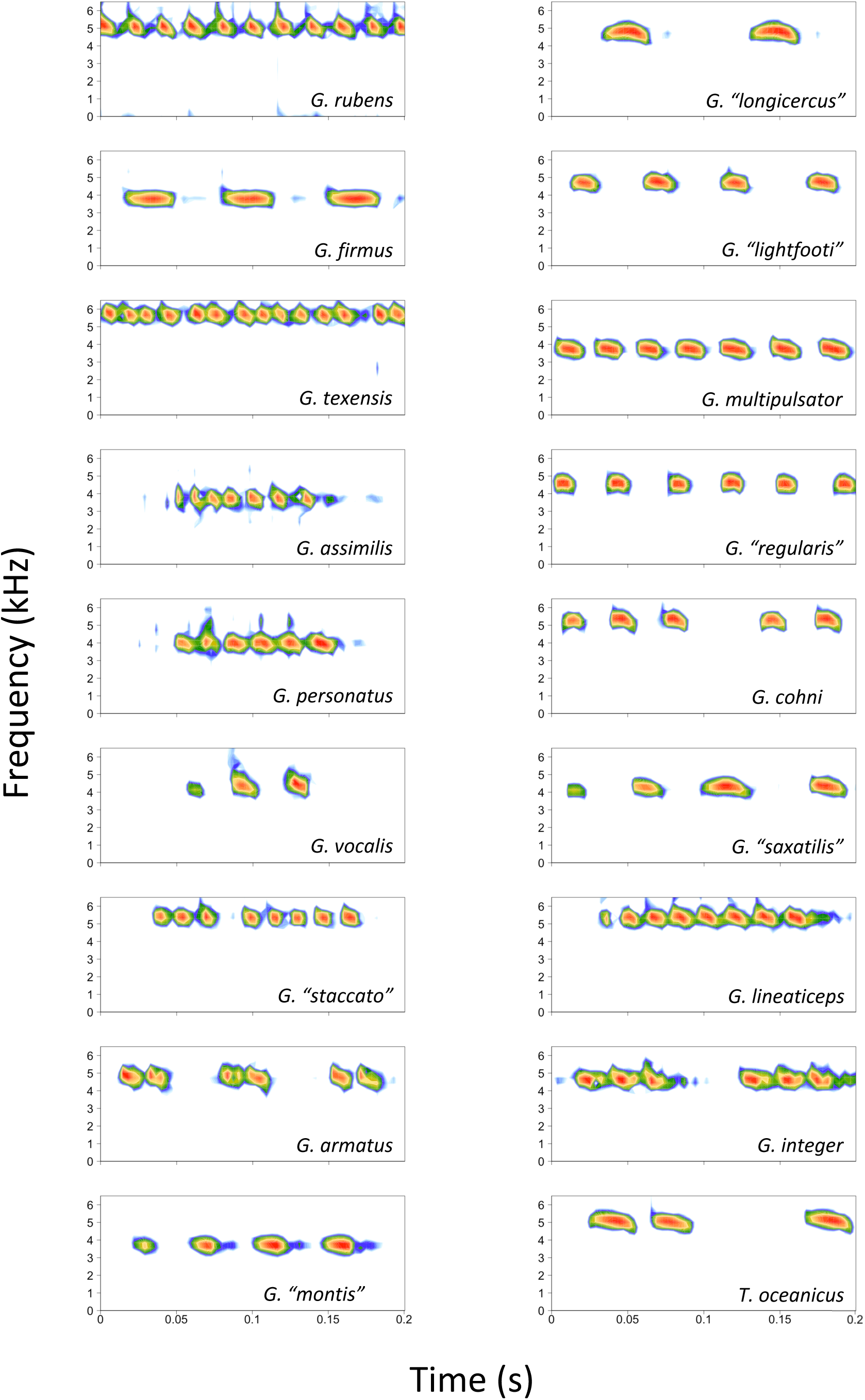
Spectrogram representations of 0.2 seconds of song from confirmed host species showing finescale

The song distance matrix shows nearly 30-fold variation among species in pairwise inter-host song distance comparisons (Fig. 6). Notably, the average distance of *T. oceanicus* song from each of the other songs was about double the average distances for the continental *Gryllus* species (7.75 versus 3.85, Z = 7.4, p < 0.0001).

**Figure 6.**
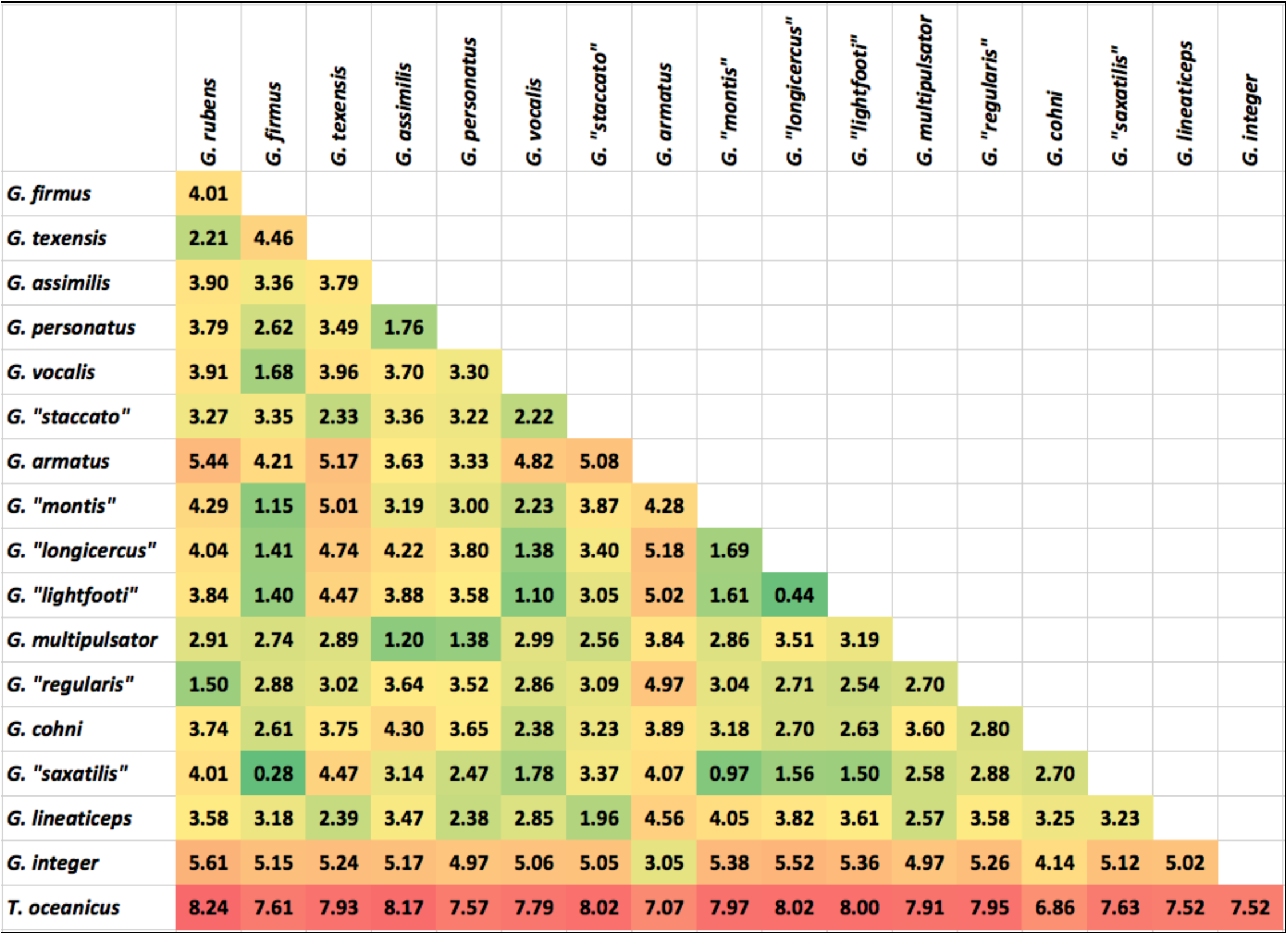
Euclidean pairwise inter-host song distances with heatmap colors indicating similar songs (green) or strongly divergent songs (red).

## Discussion

Our results suggest the following: (1) *O. ochracea* is a single widespread species with regional host specialization, not a complex of cryptic species, (2) *O. ochracea* has spread geographically into northern Mexico (Sonora) and the western USA (Arizona and California) from source populations in southern Mexico (Oaxaca) and/or the southern USA Gulf region (Florida, Texas), (3) Hawaiian flies were introduced from a western continental USA population, most likely California, potentially consisting of as few as one gravid female fly, and (4) novel song types with highly divergent song structures do not inhibit novel host exploitation. We elaborate on these results below, and discuss mechanisms of regional host song specialization.

Studies of other Tachinid groups have sometimes revealed that what was considered a single generalist species actually consists of a complex of cryptic specialist species (Smith et al., 2007, Smith et al., 2006). The regional host specialization in *O. ochracea* documented previously (Gray et al., 2007) could have been consistent with either a widespread generalist with regional host preferences or with multiple cryptic host specialists. Both the mtDNA and msat variation suggest a single species. The mtDNA sequences, although showing clear east-west geographic structure, are relatively uniform and strongly divergent from *O. depleta* and *O. lineifrons* sequences (Supplemental Materials Figure S13). The msat data clearly show that populations strongly differentiated in host song preferences can nonetheless be genetically panmictic. Perhaps the best example of this involves flies from Florida and Texas: Gray et al. (2007) showed that Florida flies preferred *G. rubens* song over *G. texensis* song nearly 2:1 and that Texas flies preferred *G. texensis* song over *G. rubens* song 6:1. Nonetheless the pairwise Fst of 0.008 for these populations (Table 3) and the DAPC (Fig. 1a) show that these two populations are genetically rather homogenous.

Both the mtDNA and msat data also inform the broader geographic history of the fly within North America. There is a clear east-west differentiation among samples, potentially consistent with isolation by distance. Moreover, the pattern of allelic variation in the msat loci (e.g. Fig. 3) suggests serial founder effects as flies colonized the western continental USA and then Hawaii. The mtDNA similarly suggests that the older fly lineages are to be found within the southeastern USA populations (Fig. 1b; Fig. S13). In this light, it is interesting to note that Florida is home to two *Gryllus* species, *G. ovisopis* and *G. cayensis,* which lack a normal calling song (Gray et al., 2018, Walker, 1974, Walker, 2001), possibly a consequence of a prolonged history of *Ormia* parasitism in that region. In contrast, there are no non-calling *Gryllus* in western North America.

The introduction of *O. ochracea* to Hawaii appears virtually certain to have been from a western North American population. The dominant mtDNA haplotype in Hawaii is also found in California and Arizona (Fig. 1b); the msat allelic variation in Hawaii is likewise a subset of the most common alleles in California and Arizona (Fig. 3). A single introduction seems likely; the levels of genetic variation in Hawaii do not preclude the possibility that the introduction could have consisted of as few as one gravid female, although it seems more plausible that multiple individuals were introduced, perhaps as pupae in soil. In other systems, experimental introductions have indicated that in some circumstances introductions of a single gravid female can nonetheless establish a persistent population (Grevstad, 1999, Fauvergue et al., 2007). Within Hawaii, our data are consistent with the spread of an introduced population among islands, rather than separate introductions on each island (Supplemental Fig. S1).

Once in Hawaii, the adoption of *T. oceanicus* as a host represents a major shift within *O. ochracea*’s repertoire of host song recognition. Quantitatively and qualitatively, *T. oceanicus* song is strikingly divergent from the songs of continental North American hosts (Figs. 4-6). Across the diversity of host songs, one could argue that the single essential song recognition feature is a dominant frequency in the 3-6 kHz range. This may be true in a strict sense, but frequency is clearly not the only song recognition feature. Multiple studies have shown that the temporal pattern of sound pulses is also important (Gray and Cade, 1999, Sakaguchi and Gray, 2011, Wagner, 1996, Wagner and Basolo, 2007, Walker, 1993). Moreover, fly populations prefer the temporal structure of their most common host species, even when dominant frequencies are similar (Gray et al., 2007). Perhaps most remarkably, Hawaiian *O. ochracea* preferred *T. oceanicus* song over the songs of ancestral host species by a large margin (12 of 13 Hawaiian flies chose *T. oceanicus* song over the songs of *G. rubens, G. texensis*, and *G. lineaticeps*).

Adoption of *T. oceanicus* as a host in Hawaii also required compatible host physiology for larval development. Although mostly confined to parasitism of adult males, *O. ochracea* can develop within a wide variety of crickets, including juveniles (Vincent and Bertram, 2009) and species not normally used as hosts (Thomson et al., 2012, Adamo et al., 1995a) including *Acheta domesticus* (Paur and Gray, 2011b, Paur and Gray, 2011a, Wineriter and Walker, 1990) which is more distantly related to *Gryllus* than is *Teleogryllus* (Gray, D.A, Weissman, D.B., Lemmon, E.M., Lemmon, A.R, unpublished data). This latitude probably results from the generalized nature of the cricket immune encapsulation response (Vinson, 1990), which is exploited by Ormiines to develop a respiratory spiracle. Given this latitude, we expect that physiological compatibility with *T. oceanicus* was unlikely to be a significant factor in terms of host suitability.

Our results suggest that host specialization in *O. ochracea* is not at odds with rapid exploitation of novel hosts, as might be expected from evolutionary theory (Raia and Fortelius, 2013, Jaenike, 1990, Kelley and Farrell, 1998). But how can highly regional host song specificity (Gray et al., 2007), even to the point of flies having song preferences for certain intra-specific song variants (Gray and Cade, 1999, Sakaguchi and Gray, 2011, Wagner, 1996, Wagner and Basolo, 2007), be compatible with flexible and rapid adoption of novel hosts? If population differentiation does not explain regional host specialization, as suggested by the results presented here, then behavioral plasticity coupled with local host learning (Paur and Gray, 2011a) may be the mechanism that enables flies to escape the ‘dead-end’ of specialization.

## Acknowledgements

We are grateful for the advice and/or assistance of Steve Bogdanowicz; Thomas Chaffee; Thomas J. Walker; David B. Weissman.

## Author Contributions

DAG, SLB, MZ, and WHC conceived of the study and collected flies; DAG performed the mtDNA sequencing; SLB and HDK performed the msat amplification and analysis; all authors contributed to the writing and editing of the manuscript.

## Data Accessibility

The COI sequence data have been deposited in GenBank with accession numbers MK522523-MK522797. Upon acceptance the msat data will be archived in Dryad.

---

**Supplemental Figure S1.**
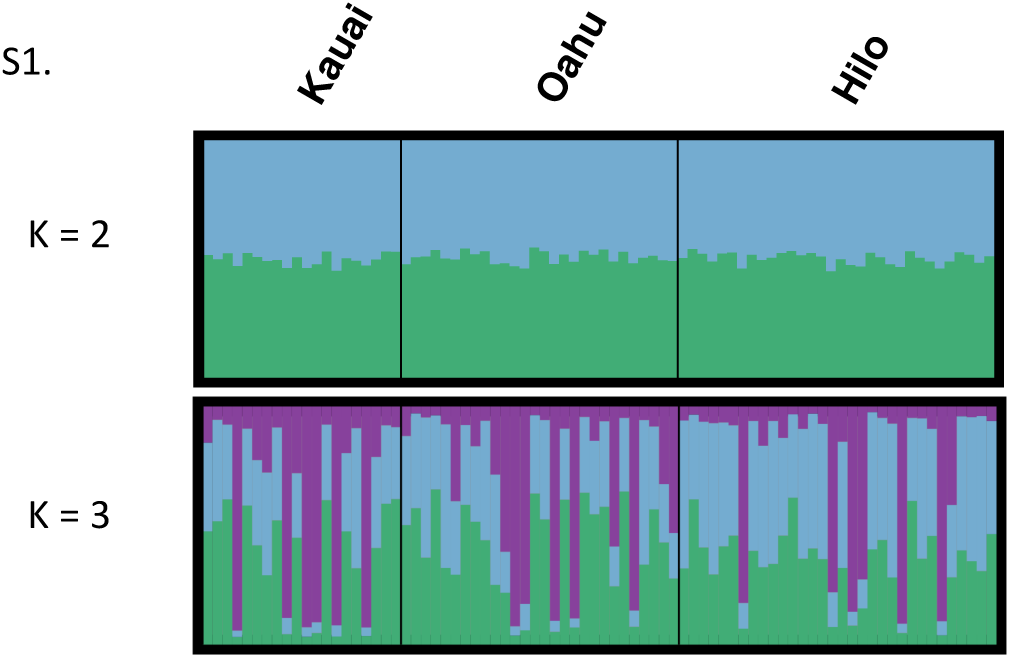
STRUCTURE plots for Hawaii flies (K=2 and K=3)

**Supplemental Figure S2.**
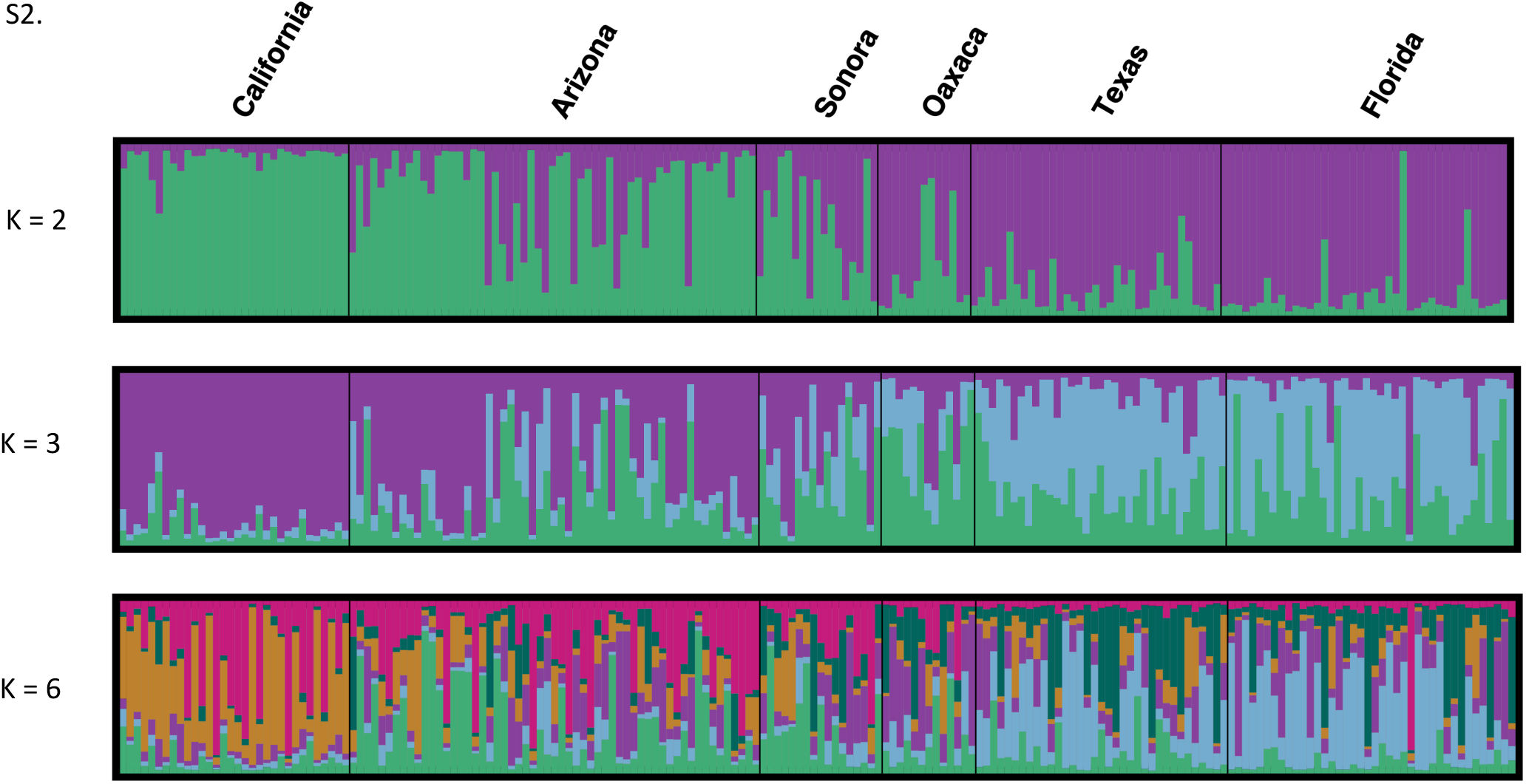
STRUCTURE plots for mainland flies (K=2, K=3, and K=6)

**Supplemental Figure S3.**
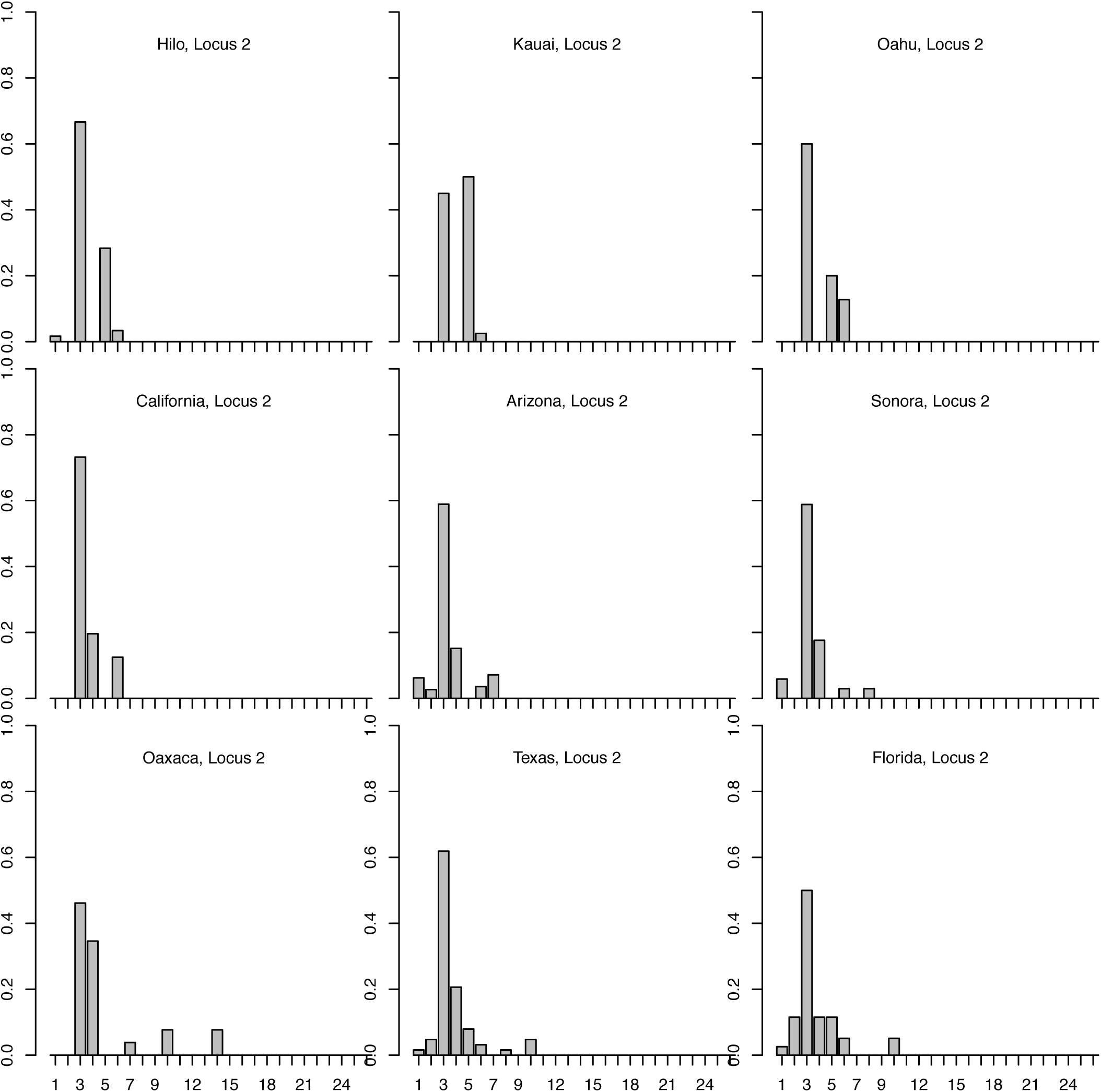
Allele frequency histograms for msat locus 2 for each population.

**Supplemental Figure S4.**
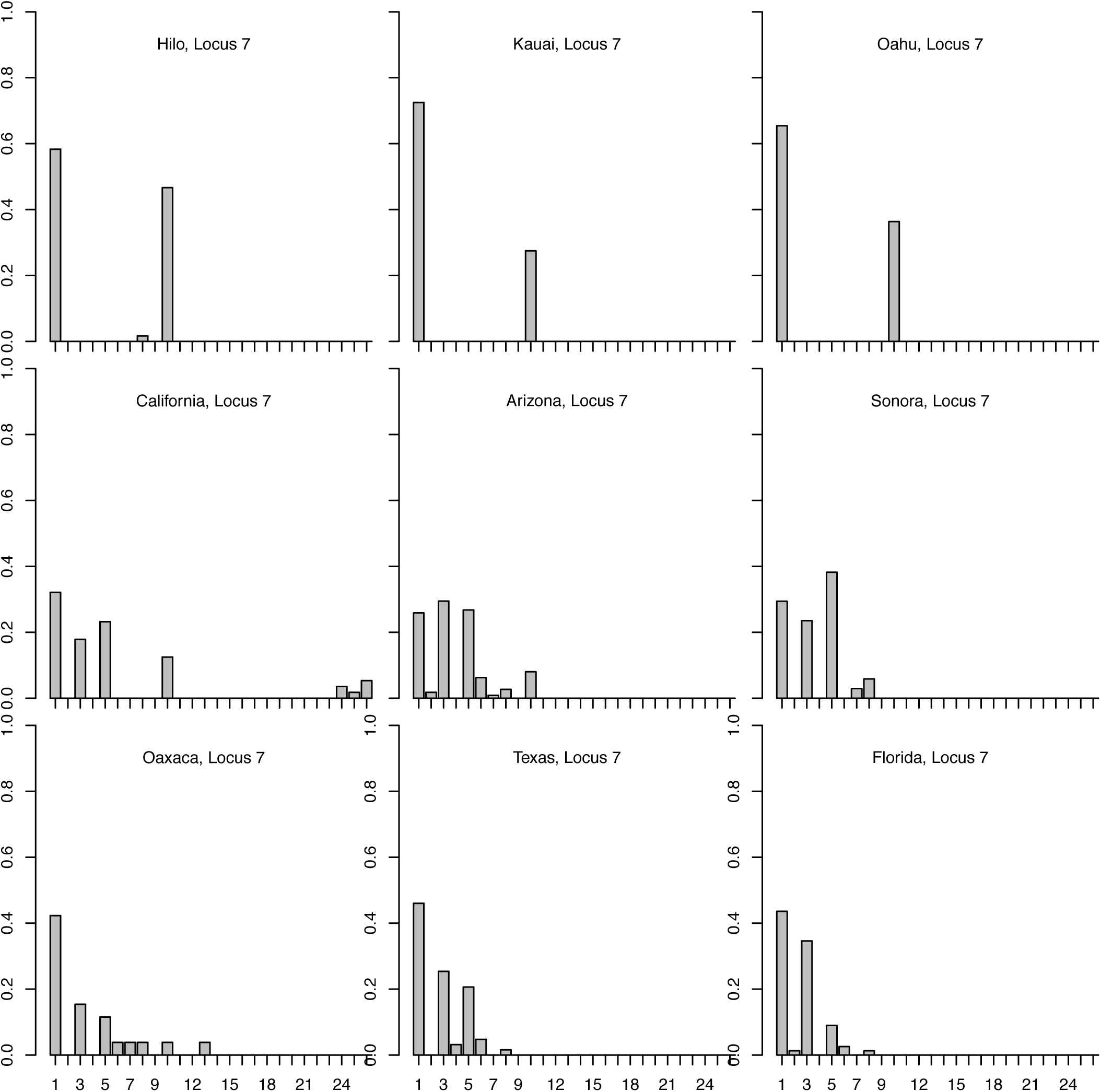
Allele frequency histograms for msat locus 7 for each population.

**Supplemental Figure S5.**
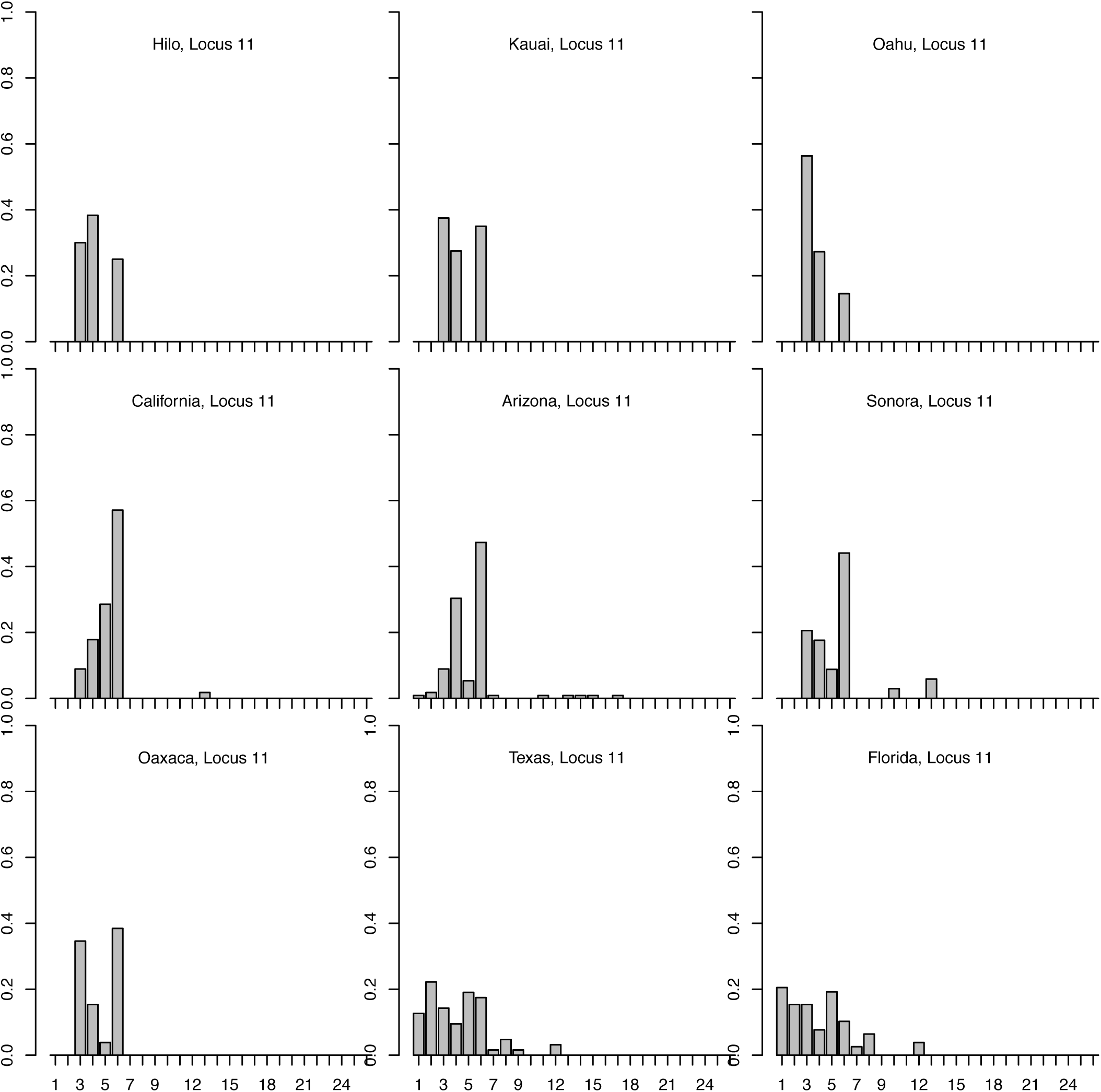
Allele frequency histograms for msat locus 11 for each population.

**Supplemental Figure S6.**
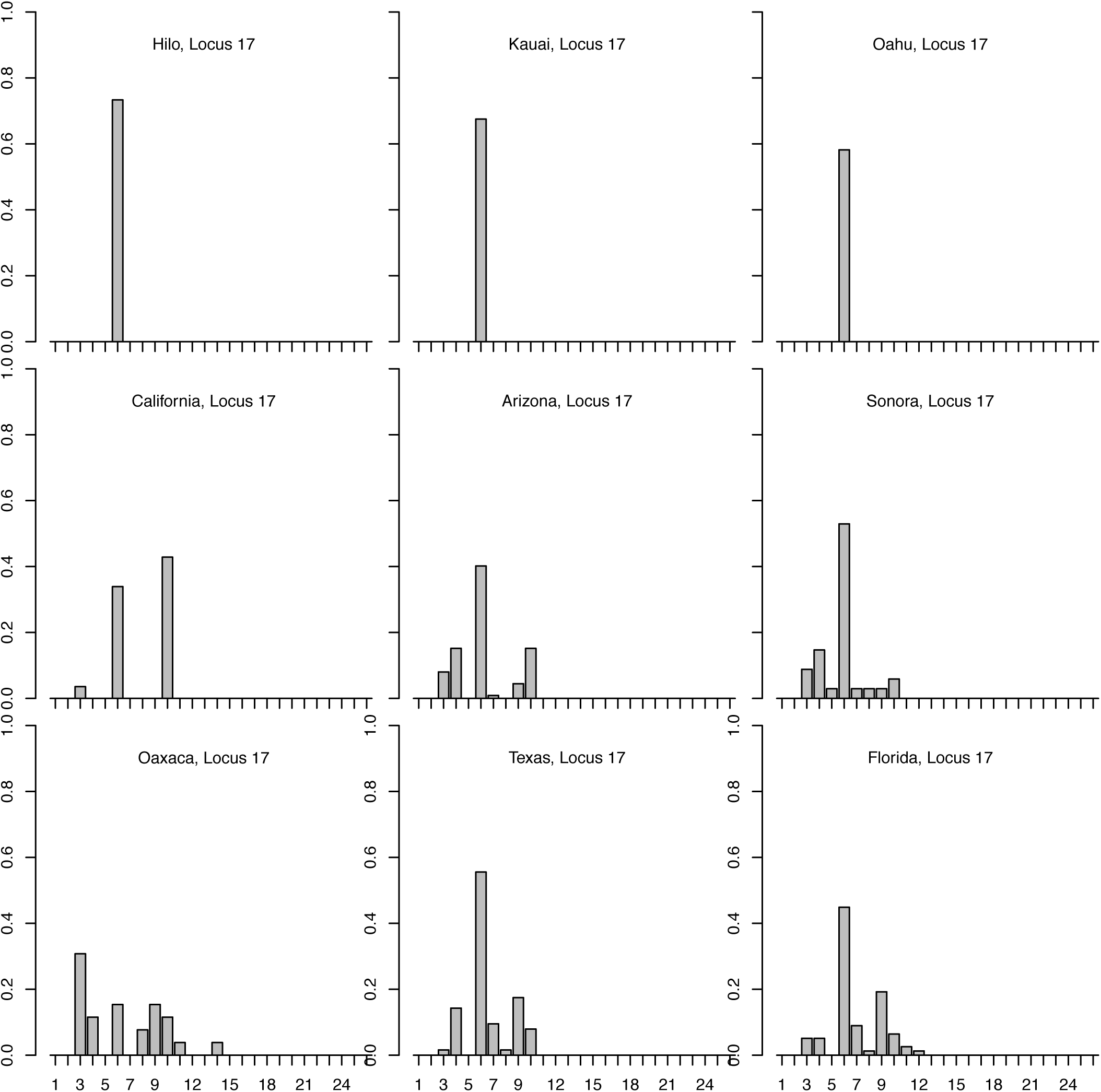
Allele frequency histograms for msat locus 17 for each population.

**Supplemental Figure S7.**
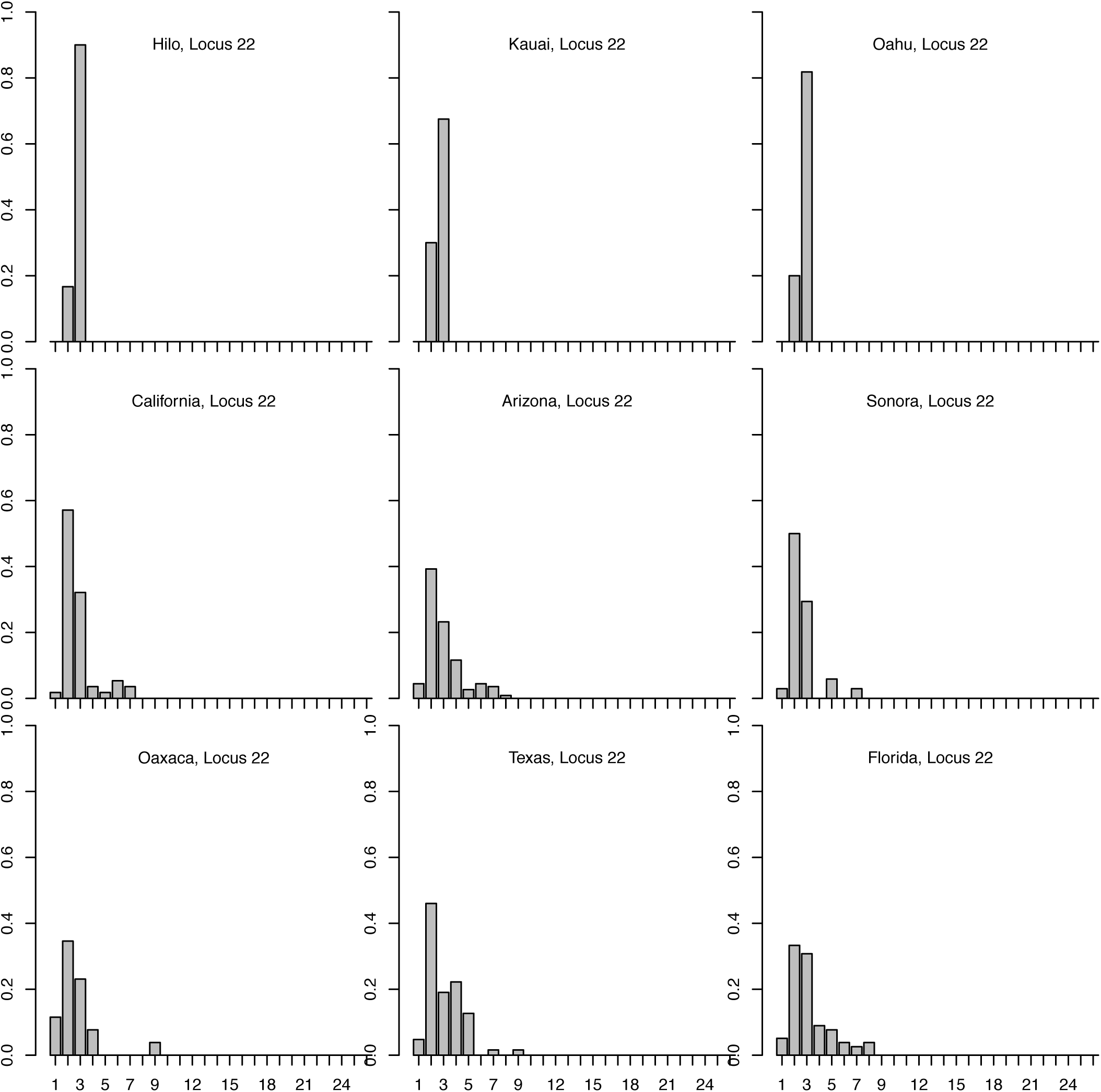
Allele frequency histograms for msat locus 22 for each population.

**Supplemental Figure S8.**
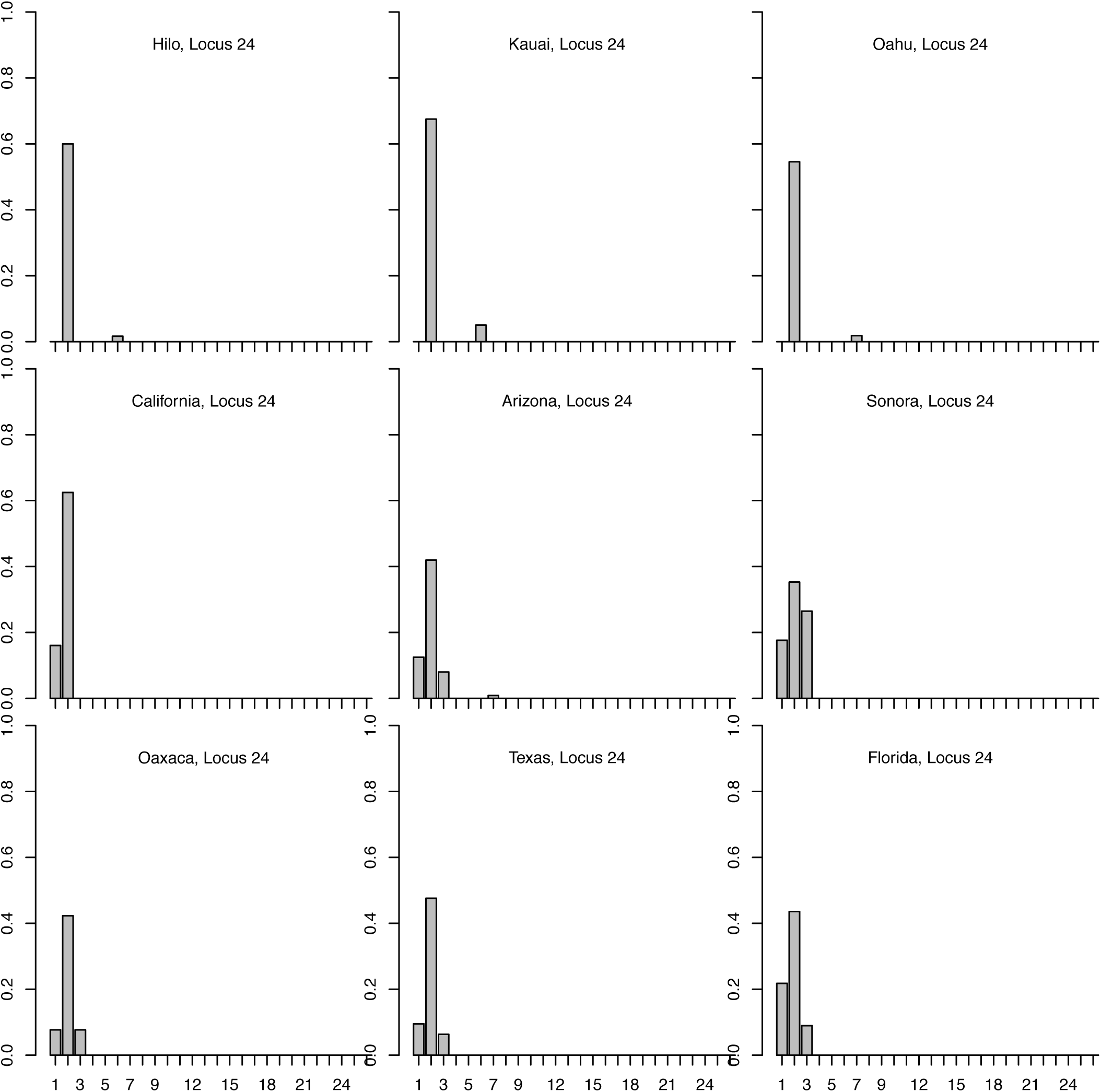
Allele frequency histograms for msat locus 24 for each population.

**Supplemental Figure S9.**
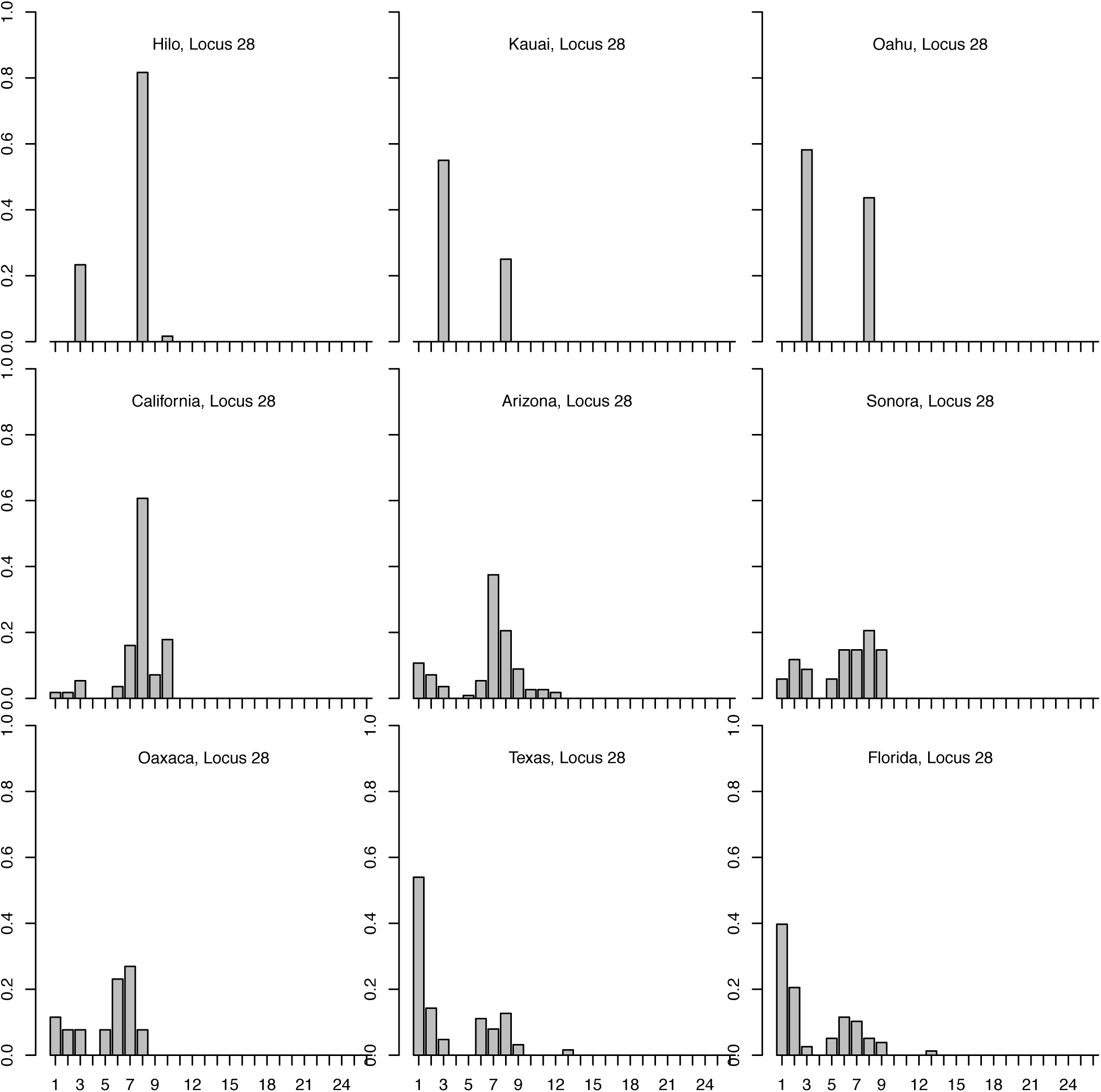
Allele frequency histograms for msat locus 28 for each population.

**Supplemental Figure S10.**
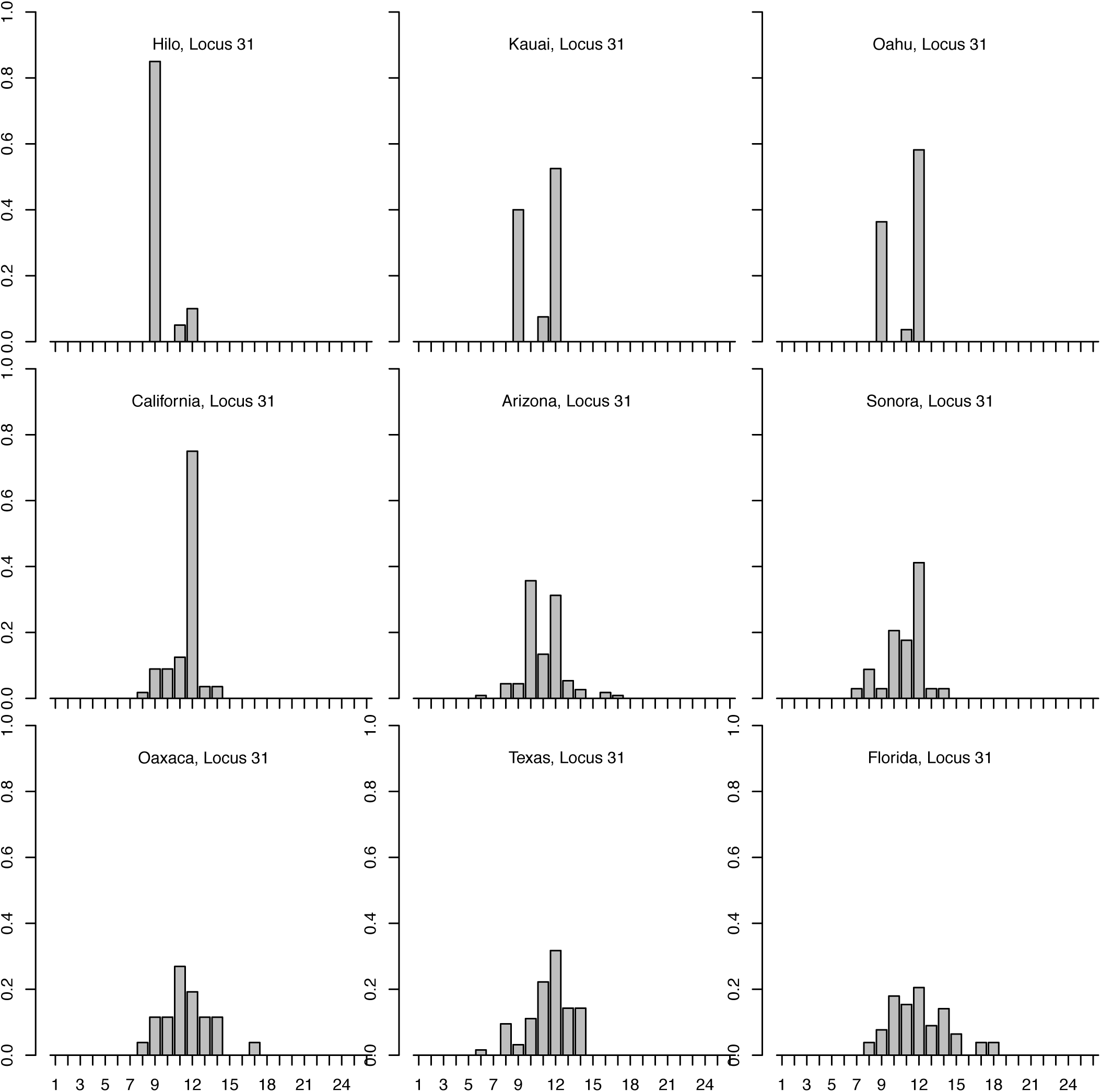
Allele frequency histograms for msat locus 31 for each population.

**Supplemental Figure S11.**
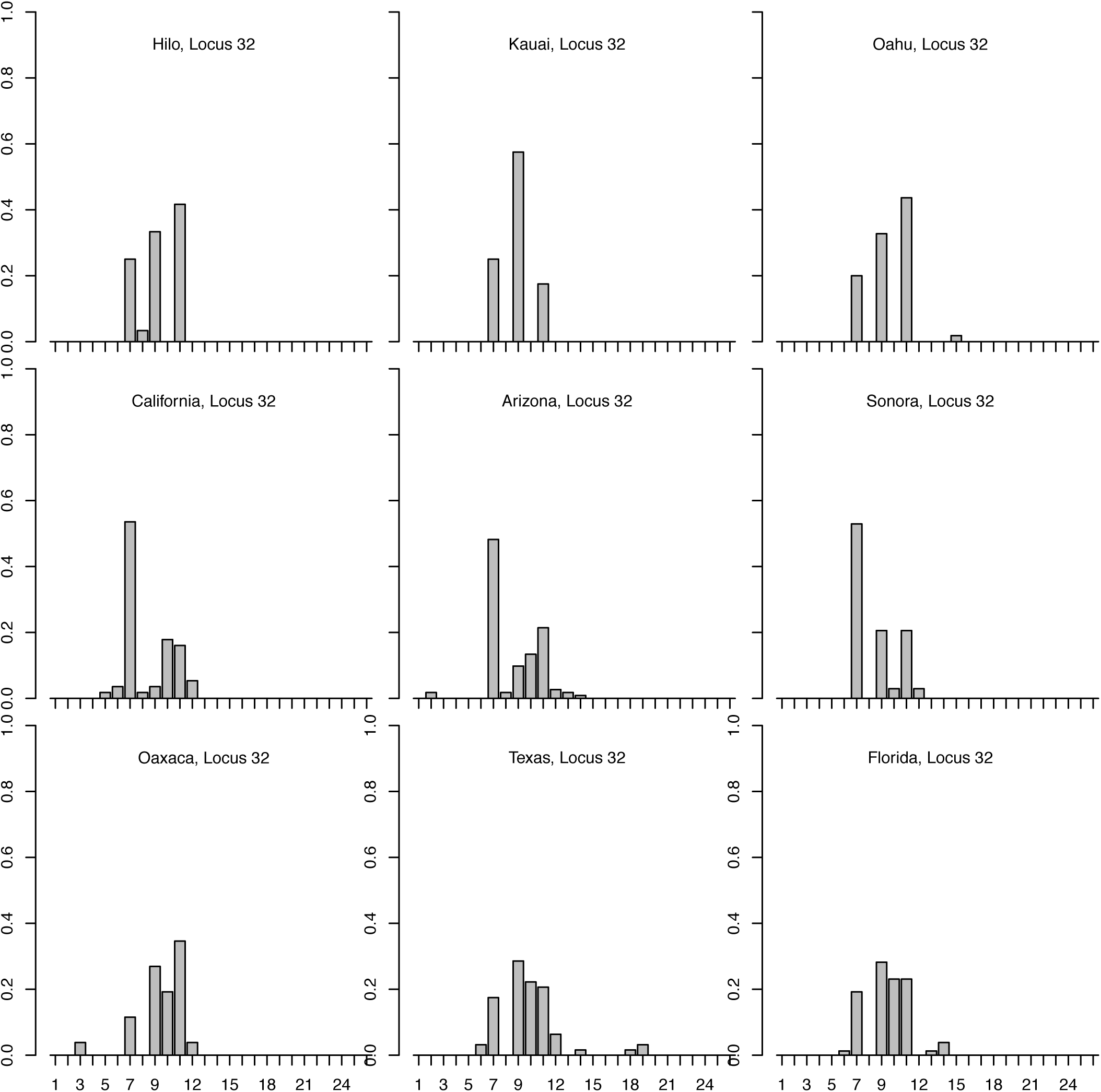
Allele frequency histograms for msat locus 32 for each population.

**Supplemental Figure S12.**
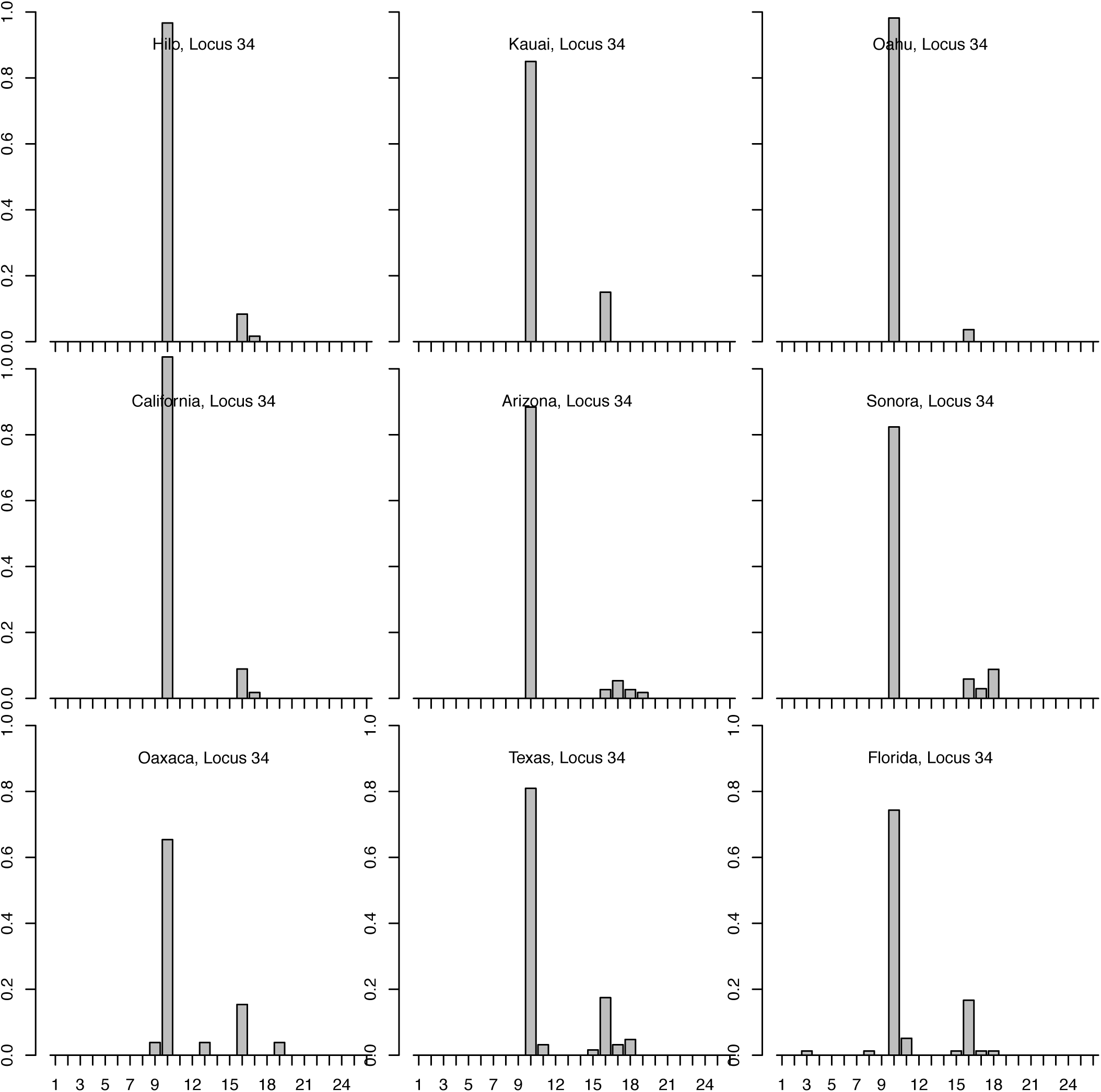
Allele frequency histograms for msat locus 34 for each population.

**Supplemental Figure S13.**
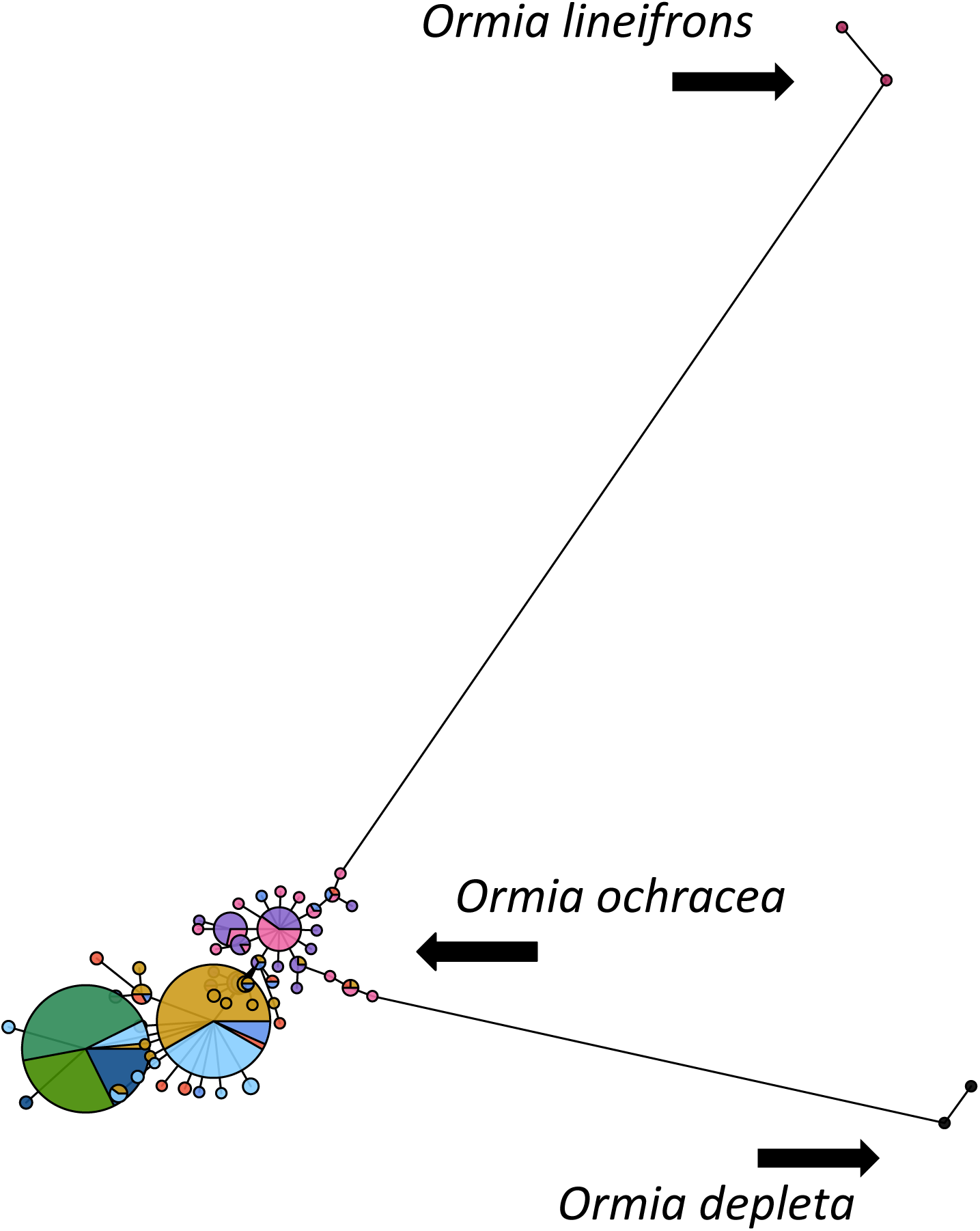
*Ormia ochracea* haplotype network with outgroups *O. lineifrons* and *O depleta* appended.

## REFERENCES

Adamo, S. A., Robert, D. & Hoy, R. R. 1995a. Effects of a tachinid parasitoid, *Ormia ochracea*, on the behaviour and reproduction of its male and female field cricket hosts (*Gryllus* spp). Journal of Insect Physiology, 41, 269–277.

Adamo, S. A., Robert, D., Perez, J. & Hoy, R. R. 1995b. The response of an insect parasitoid, *Ormia ochracea* (Tachinidae), to the uncertainty of larval success during infestation. Behavioral Ecology and Sociobiology, 36, 111–118.

Allen, G. R. 1995. The biology of the phonotactic parasitoid, *Homotrixa* sp. (Diptera: Tachinidae), and its impact on the survival of male *Sciarasaga quadrata* (Orthoptera: Tettigoniidae) in the field. Ecological Entomology, 20, 103–110.

Allen, G. R., Kamien, D., Berry, O., Byrne, p. & Hunt, J. 1999. Larviposition, host cues, and planidial behavior in the sound-locating parasitoid fly *Homotrixa alleni* (Diptera: Tachinidae). Journal of Insect Behavior, 12, 67–79.

Arnaud, P. H. 1978. Host Parasite Catalog of North American Tachinidae (Diptera). United States Department of Agriculture, Miscellaneous Publication, 1319, 1– 860.

Blackburn, T. M. & Duncan, R. P. 2001. Determinants of establishment success in introduced birds. Nature, 414, 195.

Blankers, T., Hennig, M. R. & Gray, D. A. 2015. Conservation of multivariate female preference functions and preference mechanisms in three species of trilling field crickets. J. Evol. Biol., 28, 630–641.

Cade, W. H. 1975. Acoustically orienting parasitoids: fly phonotaxis to cricket song. Science, 190, 1312–1313.

Cade, W. H. 1981. Field cricket spacing and the phonotaxis of crickets and parasitoid flies to clumped and isolated cricket songs. Zeitshrift für Tierpsychologie, 55, 365–375.

Cade, W. H., Ciceran, M. & Murray, A.-M. 1996. Temporal patterns of parasitoid fly (*Ormia ochracea*) attraction to field cricket song (*Gryllus integer*). Canadian Journal of Zoology, 74, 393–395.

Cantrell, B. 1988. The comparative morphology of the male and female postabdomen of the Australian Tachinidae (Diptera), with descriptions of some first-instar larvae and pupae. Invertebrate Systematics, 2, 81–221.

Chessel, D., Dufour, A. B. & Thioulouse, J. 2004. The ade4 package-I-One-table methods. R news, 4, 5–10.

Doherty, J. A. & Storz, M. M. 1992. Calling song and selective phonotaxis in the field crickets, *Gryllus firmus* and *G. pennsylvanicus* (Orthoptera: Gryllidae). Journal of Insect Behavior, 5, 555–569.

Edgecomb, R. S., Robert, D., Read, M. P. & Hoy, R. R. 1995. The tympanal hearing organ of a fly: phylogenetic analysis of its morphological origins. Cell and tissue research, 282, 251–268.

Evanno, G., Regnaut, S. & Goudet, J. 2005. Detecting the number of clusters of individuals using the software STRUCTURE: a simulation study. Molecular Ecology, 14, 2611–2620.

Evenhuis, N. L. 2003. The status of the cricket parasites *Ormia ochracea* and *Phasioormia pallida* in the Hawaiian Islands (Diptera: Tachinidae). Bishop Museum Occasional Papers, 74, 34–35.

Fauvergue, X., Malausa, J.-C., Giuge, L. & Courchamp, F. 2007. Invading parasitoids suffer no Allee effect: a manipulative field experiment. Ecology, 88, 2392–2403.

Gompert, Z., Jahner, J. P., Scholl, C. F., Wilson, J. S., Lucas, L. K., SoriaLCarrasco, V., Fordyce, J. A., Nice, C. C., Buerkle, C. A. & Forister, M. L. 2015. The evolution of novel host use is unlikely to be constrained by tradeLoffs or a lack of genetic variation. Molecular ecology, 24, 2777–2793.

González-Suárez, M., Bacher, S. & Jeschke, J. M. 2015. Intraspecific trait variation is correlated with establishment success of alien mammals. The American Naturalist, 185, 737–746.

Gray, D. A., Banuelos, C. M., Walker, S. E., Cade, W. H. & Zuk, M. 2007. Behavioural specialisation among populations of the acoustically-orienting parasitoid fly *Ormia ochracea* utilising different cricket species as hosts. Animal Behaviour, 73, 99–104.

Gray, D. A. & Cade, W. H. 1999. Sex, death and genetic variation: natural and sexual selection on cricket song. Proceedings of the Royal Society of London, Series B: Biological Sciences, 266, 707–709.

Gray, D. A., Gabel, E., Blankers, T. & Hennig, R. M. 2016a. Multivariate female preference tests reveal latent perceptual biases. Proc. R. Soc. B, 283, doi:10.1098/rspb.2016.1972.

Gray, D. A., Gutierrez, N. J., Chen, T. L., Gonzalez, C., Weissman, D. B. & Cole, J. A. 2016b. Species divergence in field crickets: genetics, song, ecomorphology, and pre- and postzygotic isolation. Biological Journal of the Linnaean Society, 117, 192–205.

Gray, D. A., Hormozi, S., Libby, F. R. & Cohen, R. W. 2018. Induced expression of a vestigial sexual signal. Biology Letters, 14, 20180095.

Grevstad, F. S. 1999. Experimental invasions using biological control introductions: the influence of release size on the chance of population establishment. Biological Invasions, 1, 313–323.

Hedrick, A. V. & Kortet, R. 2006. Hiding behaviour in two cricket populations that differ in predation pressure. Animal Behaviour, 72, 1111–1118.

Hedrick, A. V. & Weber, T. 1998. Variance in female responses to the fine structure of male song in the field cricket, *Gryllus integer*. Behavioral Ecology, 9, 582–591.

Hedwig, B. & Robert, D. 2014. Auditory parasitoid flies exploiting acoustic communication of insects. Insect hearing and acoustic communication. Springer.

Henne, D. C. & Johnson, S. J. 2001. Seasonal distribution and parasitism of *Scapteriscus* spp. (Orthoptera: Gryllotalpidae) in southeastern Louisiana. Florida Entomologist, 84, 209–214.

Higgins, S. I. & Richardson, D. M. 2014. Invasive plants have broader physiological niches. Proceedings of the National Academy of Sciences, 111, 10610–10614.

Izzo, A. S. & Gray, D. A. 2004. Cricket song in sympatry: species specificity of song without reproductive character displacement in *Gryllus rubens*. Annals of the Entomological Society of America, 97, 831–837.

Jaenike, J. 1990. Host specialization in phytophagous insects. Annual Review of Ecology and Systematics, 21, 243–273.

Jombart, T. 2008. adegenet: a R package for the multivariate analysis of genetic markers. Bioinformatics, 24, 1403–1405.

Jombart, T. & Ahmed, I. 2011. adegenet 1.3-1: new tools for the analysis of genome-wide SNP data. Bioinformatics, 27, 3070–3071.

Kalinowski, S. T. 2005. hpLrare 1.0: a computer program for performing rarefaction on measures of allelic richness. Molecular Ecology Notes, 5, 187–189.

Kelley, S. T. & Farrell, B. D. 1998. Is specialization a dead end? The phylogeny of host use in *Dendroctonus* bark beetles (Scolytidae). Evolution, 52, 1731–1743.

Kevan, D. K. M. 1990. Introduced grasshoppers and crickets in Micronesia. Boletin de Sanidad Vegetal, 20, 105–123.

Lehmann, G. U. C. 2003. Review of biogeography, host range and evolution of acoustic hunting in Ormiini (Insecta, Diptera, Tachinidae), parasitoids of night-calling bushcrickets and crickets (Insecta, Orthoptera, Ensifera). Zoologischer Anzeiger, 242, 107–120.

Nutting, W. L. 1953. The biology of *Euphasiopteryx brevicornis* (Townsend) (Diptera, Tachinidae), a parasite in the cone-headed grasshoppers (Orthoptera: Copiphorinae). Psyche, 60, 69–81.

Paradis, E. 2010. pegas: an R package for population genetics with an integrated– modular approach. Bioinformatics, 26, 419–420.

Parkman, J. P., Frank, J. H., Walker, T. J. & Schuster, D. J. 1996. Classical biological control of *Scapteriscus* spp. (Orthoptera: Gryllotalpidae) in Florida. Environmental Entomology, 25, 1415–1420.

Paur, J. & Gray, D. A. 2011a. Individual consistency, learning and memory in a parasitoid fly, *Ormia ochracea*. Animal Behaviour, 82, 825–830.

Paur, J. & Gray, D. A. 2011b. Seasonal dynamics and overwintering strategy of the tachinid fly, *Ormia ochracea*, in southern California. Terrestrial Arthropod Reviews, 4, 145–156.

Quicke, D. L. 2014. The braconid and ichneumonid parasitoid wasps: biology, systematics, evolution and ecology, John Wiley & Sons.

Raia, P. & Fortelius, M. 2013. Cope’s Law of the Unspecialized, Cope’s Rule, and weak directionality in evolution. Evolutionary Ecology Research, 15, 747–756.

Robert, D., Amoroso, J. & Hoy, R. R. 1992. The evolutionary convergency of hearing in a parasitoid fly and its cricket host. Science, Wash., 258, 1135–1137.

Romanuk, T. N., Zhou, Y., Brose, U., Berlow, E. L., Williams, R. J. & Martinez, N. D. 2009. Predicting invasion success in complex ecological networks. Philosophical Transactions of the Royal Society B: Biological Sciences, 364, 1743–1754.

Sabrosky, C. W. 1953a. Taxonomy and host relations of the tribe Ormiini in the western hemisphere (Diptera, Larvaevoridae). Proceedings of the Entomological Society of Washington, 55, 167–183.

Sabrosky, C. W. 1953b. Taxonomy and host relations of the tribe Ormiini in the western hemisphere, II (Diptera, Larvaevoridae). Proceedings of the Entomological Society of Washington, 55, 289–305.

Sakaguchi, K. M. & Gray, D. A. 2011. Host song selection by an acoustically-orienting parasitoid fly exploiting a multi-species assemblage of cricket hosts. Animal Behaviour, 81, 851–858.

Shapiro, L. 1995. Parasitism of *Orchelimum* katydids (Orthoptera: Tittigoniidae) by *Ormia lineifrons* (Diptera: Tachinidae). Florida Entomologist, 78, 615–616.

Simon, C., Frati, F., Beckenbach, A., Crespi, B., Liu, H. & Flook, P. 1994. Evolution, weighting, and phylogenetic utility of mitochondrial gene sequences and a compilation of conserved polymerase chain reaction primers. Annals of the Entomological Society of America, 87, 651–701.

Smith, M. A., Rodriguez, J. J., Whitfield, J. B., Deans, A. R., Janzen, D. H., Hallwachs, W. & Hebert, P. D. 2008. Extreme diversity of tropical parasitoid wasps exposed by iterative integration of natural history, DNA barcoding, morphology, and collections. Proceedings of the National Academy of Sciences, 105, 12359–12364.

Smith, M. A., Wood, D. M., Janzen, D. H., Hallwachs, W. & Hebert, P. D. 2007. DNA barcodes affirm that 16 species of apparently generalist tropical parasitoid flies (Diptera, Tachinidae) are not all generalists. Proceedings of the National Academy of Sciences, U S A, 104, 4967–72.

Smith, M. A., Woodley, N. E., Janzen, D. H., Hallwachs, W. & Hebert, P. D. 2006. DNA barcodes reveal cryptic host-specificity within the presumed polyphagous members of a genus of parasitoid flies (Diptera: Tachinidae). Proceedings of the National Academy of Sciences, U S A, 103, 3657–62.

Snyder, W. E. & Evans, E. W. 2006. Ecological effects of invasive arthropod generalist predators. Annu. Rev. Ecol. Evol. Syst., 37, 95–122.

Stireman, J. 2005. The evolution of generalization? Parasitoid flies and the perils of inferring host range evolution from phylogenies. Journal of evolutionary biology, 18, 325–336.

Stireman, J. O., O’Hara, J. E. & Wood, D. M. 2006. Tachinidae: Evolution, behavior, and ecology. Annual Review of Entomology, 51, 525–555.

Thomson, I. R., Vincent, C. M. & Bertram, S. 2012. Success of the parasitoid fly *Ormia ochracea* (Diptera: Tachinidae) on natural and unnatural cricket hosts. Florida Entomologist, 95, 43–48.

Tinghitella, R., Zuk, M., Beveridge, M. & Simmons, L. 2011. Island hopping introduces Polynesian field crickets to novel environments, genetic bottlenecks and rapid evolution. Journal of evolutionary biology, 24, 1199–1211.

Tschorsnig, H.-P. 2017. Preliminary host catalog of Palearctic Tachinidae (Diptera) [Online]. Available: http://www.nadsdiptera.org/Tach/WorldTachs/CatPalHosts/Cat_Pal_tach_hosts_Ver1.pdf [Accessed June 28, 2018].

Vamosi, J. C., Armbruster, W. S. & Renner, S. S. 2014. Evolutionary ecology of specialization: insights from phylogenetic analysis. 281, 2014.2004.

Vargas, R. I., Leblanc, L., Putoa, R. & Eitam, A. 2007. Impact of introduction of *Bactrocera dorsalis* (Diptera: Tephritidae) and classical biological control releases of *Fopius arisanus* (Hymenoptera: Braconidae) on economically important fruit flies in French Polynesia. Journal of Economic Entomology, 100, 670–679.

Vélez, M. J. & Brockmann, H. J. 2006. Seasonal variation in selection on male calling song in the field cricket, *Gryllus rubens*. Animal Behaviour, 72, 439–448.

Vincent, C. M. & Bertram, S. M. 2009. The parasitoid fly *Ormia ochracea* (Diptera: Tachinidae) can use juvenile crickets as hosts. Florida Entomologist, 92, 598–600.

Vinson, S. B. 1990. How parasitoids deal with the immune system of their host: an overview. Archs Insect Biochem Physiol, 13, 3–27.

Wagner, W. E. & Basolo, A. L. 2007. Host preferences in a phonotactic parasitoid of field crickets: the relative importance of host song characters. Ecological Entomology, 32, 478–484.

Wagner, W. E., Jr. 1996. Convergent song preferences between female field crickets and acoustically orienting parasitoid flies. Behavioral Ecology, 7, 279–285.

Walker, T. J. 1974. *Gryllus ovisopis* N. SP.: a taciturn cricket with a life cycle suggesting allochronic speciation. Florida Entomologist, 57, 13–22.

Walker, T. J. 1989. A live trap for monitoring *Euphasiopteryx* and tests with *E. ochracea* [Diptera: Tachinidae]. Florida Entomologist, 72, 314–319.

Walker, T. J. 1993. Phonotaxis in female *Ormia ochracea* (Diptera: Tachinidae), a parasitoid of field crickets. Journal of Insect Behavior, 6, 389–410.

Walker, T. J. 2001. Gryllus cayensis n.sp. (Orthoptera: Gryllidae), a taciturn wood cricket extirpated from the Florida Keys: songs, ecology and hybrids. Florida Entomologist, 84, 700–705.

Walker, T. J. & Wineriter, S. A. 1991. Hosts of a phonotactic parasitoid and levels of parasitism (Diptera: Tachinidae: *Ormia ochracea*). Florida Entomologist, 74, 554–559.

Weir, B. S. & Cockerham, C. C. 1984. Estimating FLstatistics for the analysis of population structure. Evolution, 38, 1358–1370.

Weissman, D. B., Rentz, D. C. F., Alexander, R. D. & Loher, W. 1980. Field crickets (*Gryllus* and *Acheta*) of California and Baja California, Mexico (Orthoptera: Gryllidae: Gryllinae). Trans. Am. Entomol. Soc., 106, 327–356.

Weissman, D. B., Walker, T. J. & Gray, D. A. 2009. The Jamaican field cricket *Gryllus assimilis* and two new sister species (Orthoptera: Gryllidae). Annals of the Entomological Society of America, 102, 367–380.

Wineriter, S. A. & Walker, T. J. 1990. Rearing phonotactic parasitoid flies (Diptera: Tachinidae, Ormiiini, *Ormia* spp.). Entomophaga, 35, 621–632.

Zuk, M., Simmons, L. W. & Cupp, L. 1993. Calling characteristics of parasitized and unparasitized populations of the field cricket *Teleogryllus oceanicus*. Behavioral Ecology and Sociobiology, 33, 339–343.

Zuk, M., Simmons, L. W. & Rotenberry, J. T. 1995. Acoustically-orienting parasitoids in calling and silent males of the field cricket *Teleogryllus oceanicus*. Ecological Entomology, 20, 380–383.

